# Stoichiometry-dependent gating and structural pharmacology of cardiac GIRK1/4 channels

**DOI:** 10.64898/2026.10.06.756879

**Authors:** Liang Wang, Takeharu Kawano, Ana Santa-Cruz, Nousheen Parvaiz, Andrew Zorn, Yongcheng Lu, Mehreen Zaka, Nihal Aly, James Pentikis, Evangelos Dadiotis, Kim W. Chan, Meng Cui, Leigh D. Plant, Diomedes E. Logothetis, Georgios Skiniotis

## Abstract

Cardiac G protein-gated inwardly rectifying potassium-K^+^ (GIRK) channels, formed by GIRK1/4 heterotetramers or GIRK4 homotetramers, are crucial regulators of heart rate, and their dysfunction is associated with atrial fibrillation. Despite their therapeutic potential as anti-arrhythmic targets, development of selective GIRK modulators has been limited by an incomplete understanding of their structures, gating mechanisms, and pharmacology. Here, we report high-resolution structures of the GIRK4 homotetramer and of GIRK1/4 heterotetramers, revealing that they form in unexpected 1:3 and 2:2 assembly stoichiometries that we corroborate using cell-based single molecule microscopy. Electrophysiological assays and molecular dynamics simulations further show that incorporation of one or two GIRK1 subunits into GIRK4-containing channels progressively opens the helix bundle crossing (HBC) gate and increases basal channel activity. Additional structures of GIRK1/4 complexes with ML297, a small-molecule ligand selective for GIRK1-containing channels, reveal distinct GIRK1-centered intersubunit binding environments. Remarkably, ML297 inhibits GIRK1/4 channels with a 1:3 stoichiometry but potentiates channels with a 2:2 stoichiometry, establishing a direct link between subunit composition, ligand-binding environment, and pharmacological efficacy. Together, these findings provide a structural and mechanistic framework for the rational design of ligands with enhanced selectivity among cardiac and neuronal GIRK channel heterotetramers.

## Introduction

Much of the progress made in identifying the G protein-gated inwardly rectifying potassium-K^+^ (GIRK) channels and elucidating their regulation by G proteins arose from efforts to understand vagally induced hyperpolarization of cardiac muscle^1–13^. Early studies established that acetylcholine and adenosine, acting through M2-muscarinic and adenosine G-protein-coupled receptors (GPCRs), respectively, activate heterotrimeric G proteins whose βγ subunits directly open GIRK channels in cardiac myocytes^13^. The resulting potassium flux generates the acetylcholine-activated current (I_K-ACh_), hyperpolarizing atrial tissue, slowing heart rate and regulating heart rate variability (HRV)^14^.

In the heart, GIRK channel activity arises from the expression of GIRK1^15^ (Kir3.1 or G1) and GIRK4^16^ (Kir3.4 or G4) subunits, which assemble into GIRK4 homotetramers and GIRK1/4 heterotetramers^16^. Genetic ablation of these channels markedly reduces the vagal chronotropic response and HRV^17,18^, while both loss- and gain-of-function mutations are linked to arrhythmias, including age-dependent atrial fibrillation, the most common form of arrhythmia in humans^14,19–21^. Despite this strong genetic and physiological evidence implicating cardiac GIRKs as therapeutic targets, efforts to develop selective modulators have been hampered by incomplete understanding of their subunit composition, gating mechanisms, and pharmacology.

Structural insight into GIRK channels has come primarily from studies of GIRK2^22–24^, which revealed a symmetric homotetramer comprising a transmembrane domain (TMD) and a large cytoplasmic domain (CTD) that together form two principal K^+^ gating constrictions: the helix bundle crossing (HBC) and the G-loop gate. Channel activation critically depends on phosphatidylinositol 4,5-bisphosphate (PI(4,5)P_2_ or PIP_2_), which stabilizes conductive conformations and enables further activation by Na^+^ and Gβγ subunits^25–27^ through mechanisms thought to be conserved across the GIRK family. However, how heteromeric assembly with GIRK1 remodels channel architecture and key structural elements governing gating and ion conduction has remained unresolved.

Biochemical and functional studies have suggested an alternating 2:2 arrangement of GIRK1 and GIRK4 subunits^28,29^, yet direct structural evidence has been lacking. This uncertainty has important pharmacological implications. ML297^30^, a small-molecule GIRK activator with pronounced selectivity for neuronal GIRK1/2 channels, also activates GIRK1/4 channels but fails to stimulate GIRK2 or GIRK4 homotetramers, implying a binding site that depends on GIRK1 incorporation. How ML297 engages heteromeric GIRKs, and how subunit compositions influence its functional effects, remain unclear.

Here, we report six high-resolution cryo-electron microscopy structures of GIRK4-containing channels, including the GIRK4 homotetramer, GIRK1/4 heterotetramers with distinct functional stoichiometries, and GIRK1/4 complexes bound to ML297. By integrating structural analysis with electrophysiology, single-molecule microscopy, mutagenesis and molecular dynamics simulations, we show that progressive incorporation of GIRK1 alters HBC gating and creates distinct intersubunit environments that determine the pharmacological responses to ML297. These findings establish a direct structural link between subunit stoichiometry, channel gating, and ligand action, providing a framework for the development of subtype- and tissue-selective GIRK modulators.

## Results

### Asymmetric stoichiometries of human GIRK1/4

To isolate human GIRK1/4 heterotetramer for structural studies, we co-expressed differentially tagged GIRK1 and GIRK4 linked by a self-cleaving P2A peptide and isolated the complex by tandem affinity purification and size exclusion chromatography (Extended Data Fig. 1a,b). Using single-particle cryo-electron microscopy (cryo-EM), we determined two structures in the absence of activator and three in the presence of ML297.

These structures revealed two distinct stoichiometries: 1:3 (GIRK1/4^1:3^) and 2:2 (GIRK1/4^2:2^) (Fig. 1a,b and Extended Data Fig. 1c). Both assemblies differed from previously presumed arrangements: the 1:3 assembly was entirely unexpected (Fig. 1a,c), whereas the anticipated 2:2 stoichiometry did not adopt the alternating configuration proposed previously^28^ but instead formed a parallel 2:2 arrangement (Fig. 1b). Subunit assignments were further supported by well-resolved side-chain densities at positions that distinguish GIRK1 from GIRK4. In both stoichiometries, GIRK1- and GIRK4-specific residues could be assigned unambiguously throughout the transmembrane domain and extracellular vestibule, confirming the identity and register of each subunit in the 1:3 and parallel 2:2 assemblies (Extended Data Fig. 1d).

**Fig. 1.**
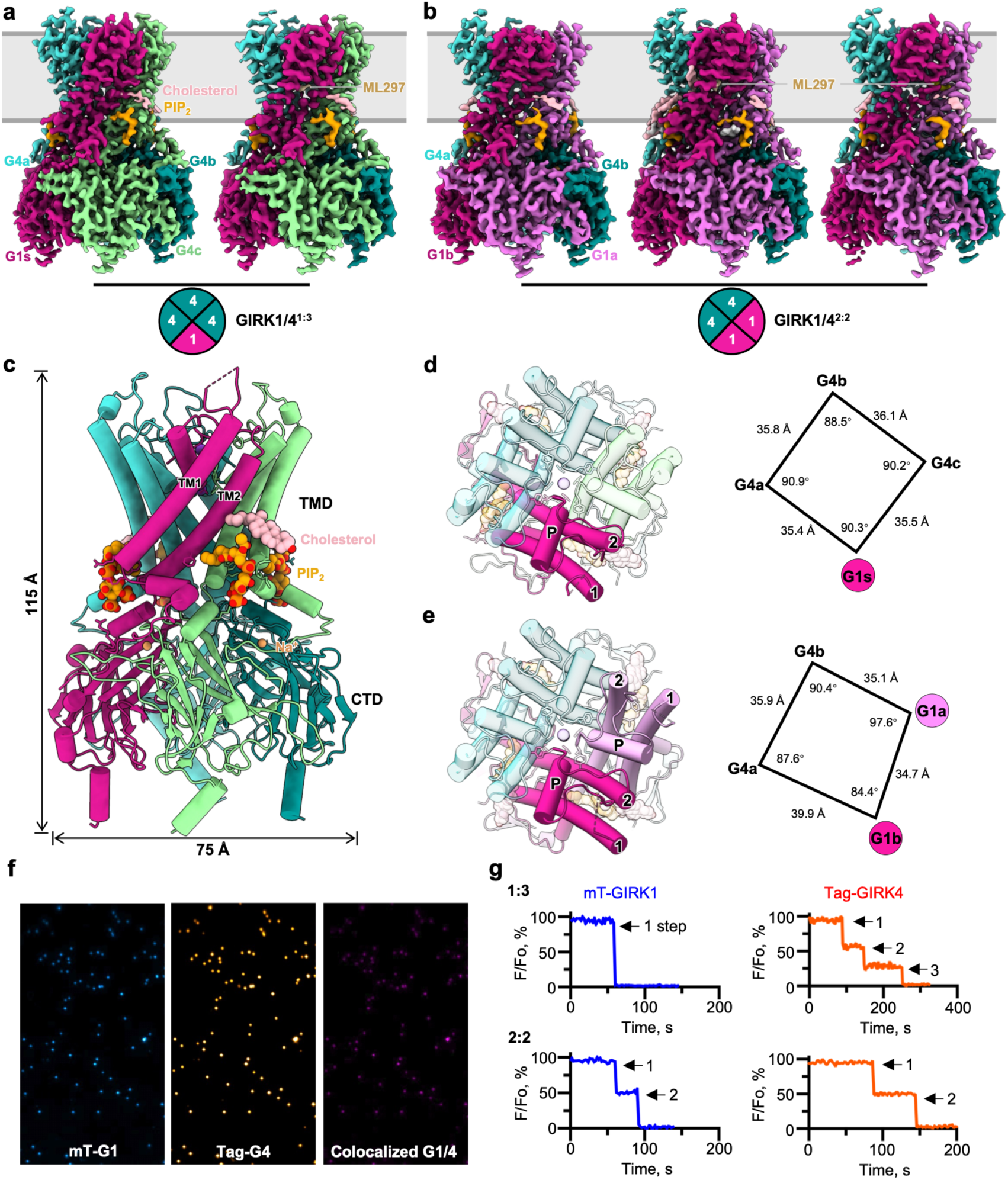
Structures and stoichiometries of GIRK1/4 heterotetramers. **a**,**b**, Cryo-EM maps of GIRK1/4 heterotetramers with 1:3 (GIRK1/4^1:3^) (**a**) and 2:2 (GIRK1/4^2:2^) (**b**) stoichiometries in complex with PIP2, cholesterol, and ML297. Each subunit is colored distinctly, and the approximate membrane boundaries are indicated in grey. Insets show schematic subunit arrangements. **c**, Side view of the atomic model of GIRK1/4^1:3^. **d**,**e**, Top-view of GIRK1/4^1:3^ (**d**) and GIRK1/4^2:2^ (**e**) showing the asymmetric subunit organization. Labels “P”, “1”, and “2” correspond to the pore helix, TM1, and TM2, respectively. Right, inter-subunit distances and angles were measured between the apices of TM1 (Cα atoms of Tyr107 in GIRK1 and Tyr113 in GIRK4), marking the TM-tip positions that define the tetrameric geometry for GIRK1/4^1:3^ (**d**) and GIRK1/4^2:2^ (**e**). **f**, Representative TIRF micrographs from a *Xenopus* oocyte expressing mTFP1-tagged GIRK1 (mT-G1) and TagRPT-T-tagged GIRK4 (Tag-G4) at a 1:1 ratio. Individual fluorescent particles are visualized. Colocalized particles are shown in orange. **g**, *Upper,* Representative photobleaching records from a colocalized particle with a single GIRK1 (blue) and three GIRK4 (red) subunits, indicating a GIRK1/4^1:3^ stoichiometry. *Lower*, Representative colocalized particle comprised of two GIRK1 and two GIRK4 subunits (GIRK1/4^2:2^ stoichiometry).

For the 1:3 assembly, we obtained an activator-free map at 2.3 Å resolution and an ML297-bound 2.8 Å map in the presence of PIP_2_, cholesterol and Na^+^ ions (Extended Data Figs. 2-4). Subunits are denoted clockwise as G1s for the single GIRK1 and G4a, G4b, and G4c for the three GIRK4 subunits, yielding the G1s-G4a-G4b-G4c configuration (Fig. 1a,d). For the 2:2 assembly, we obtained an activator-free map at 2.5 Å in complex with PIP_2_, cholesterol and one Na^+^ ion bound to the G4b subunit (Extended Data Figs. 2 and 3). Two additional ML297-bound maps at 2.4 Å and 2.8 Å, revealed ML297 binding at different subunits within the complex (Extended Data Figs 3 and 4; Fig. 1b). In the GIRK1/4^2:2^ assembly, the adjacent GIRK1 subunits are designated G1a and G1b and the adjacent GIRK4 subunits G4a and G4b (Fig. 1b,e).

The two GIRK1/4 heterotetramers adopt an overall architecture similar to GIRK2^22–24,31^, forming a channel approximately 115 Å in height and 75 Å in width, with four subunits assembling into a transmembrane domain (TMD) and a cytoplasmic domain (CTD) that dock together in the presence of PIP_2_ (Fig. 1c). However, both stoichiometries lack internal symmetry. Unlike the symmetric GIRK2 homotetramers, the TM1-tip measurements reveal asymmetric backbones: in GIRK1/4^1:3^, the angle measured for G1s differ from that of the opposing G4b subunit by ∼2° (Fig. 1d), whereas in the GIRK1/4^2:2^ complex, the largest angular difference between the opposing G1a– G4a subunits reaches ∼10° (Fig. 1e). Thus, incorporation and arrangement of GIRK1 introduce global geometric asymmetry in both GIRK1/4 assemblies.

Sequence alignment analysis shows 43.7% identity and 54.1% similarity between GIRK1 and GIRK4, with substantial differences located in the unresolved N- and C-terminal segments (residues 1-42 and 369-501 in GIRK1; 1-49 and 375-419 in GIRK4), TM1, and the turret region (Extended Data Fig. 5). Structural comparison of the GIRK1 and GIRK4 subunits within the heterotetramers reveals three prominent differences: (i) the GIRK1 turret is less well resolved, suggesting greater flexibility; (ii) the GIRK1 TM2 helix bends in the opposite direction of that in GIRK4; and (iii) the GIRK1 C-terminal helix tilts closer to 90° relative to the membrane (Extended Data Fig. 5a). Thus, GIRK1 and GIRK4 adopt distinct conformations within the asymmetric heterotetramers.

To determine whether the GIRK1/4 stoichiometries observed by cryo-EM are also formed at the plasma membrane, we performed single-molecule photobleaching using total internal reflection fluorescence (TIRF) microscopy in *Xenopus* oocytes expressing C-terminally tagged GIRK1-mTFP1 (GIRK1-TFP) and GIRK4-TagRFP-T (GIRK4-RFP). Colocalized particles showed stepwise photobleaching in both fluorescent channels, consistent with GIRK1/4 heteromers containing either one or two GIRK1 subunits (Fig. 1f,g). Across different GIRK1:GIRK4 cRNA ratios, the two subunits remained robustly colocalized at the plasma membrane (Extended Data Fig. 6a,b). To account for incomplete fluorophore detection, we further fitted the joint GIRK1 and GIRK4 bleaching-step distributions using a binomial photobleaching model incorporating three possible heteromeric stoichiometries: 1:3, 2:2 and 3:1. Across 3,575 colocalized heteromers from five cRNA ratios, the model identified a predominant 1:3 population, with a weighted average of 65%, and a substantial 2:2 population of 29%. A minor 3:1 population was also detected at approximately 6% (Extended Data Fig. 6b,c and Supplementary Table 1).

Increasing the GIRK1 cRNA fraction shifted the population toward 2:2 channels, which reached 42% in the 9:1 condition, whereas lower GIRK1 expression levels favored the 1:3 assembly (Extended Data Fig. 6b,c). GIRK4 expressed alone followed a tetrameric distribution, validating the photobleaching analysis using a known tetramer, whereas GIRK1 expressed alone produced few detectable membrane particles, consistent with inefficient surface expression of GIRK1 in the absence of a partner subunit (Extended Data Fig. 6c and Supplementary Table 1). Cryo-EM particle classification independently showed that approximately 77% of particles corresponded to the 1:3 assembly and approximately 23% corresponded to the 2:2 assembly (Extended Data Fig. 6d). Thus, GIRK1/4 forms coexisting asymmetric heterotetramers in both expression systems, with the 1:3 assembly representing the dominant species.

### GIRK1 incorporation alters pore architecture and ion conduction

To assess the functional consequences of subunit stoichiometry, we generated concatenated constructs corresponding to defined GIRK1/4^1:3^ and GIRK1/4^2:2^ assemblies and expressed them in HEK293T cells. Whole-cell patch-clamp recordings revealed robust currents for both heterotetramers as well as for the GIRK4 homotetramer (Fig. 2a). Quantification of current density revealed a clear rank order of basal activity: GIRK4 homotetramers exhibited the smallest currents, GIRK1/4^1:3^ channels displayed intermediate currents, and GIRK1/4^2:2^ channels exhibited the largest currents. All constructs retained appropriate K⁺ selectivity and inward rectification, as confirmed by their responses to high extracellular K⁺ and Ba^2^⁺.

**Fig. 2.**
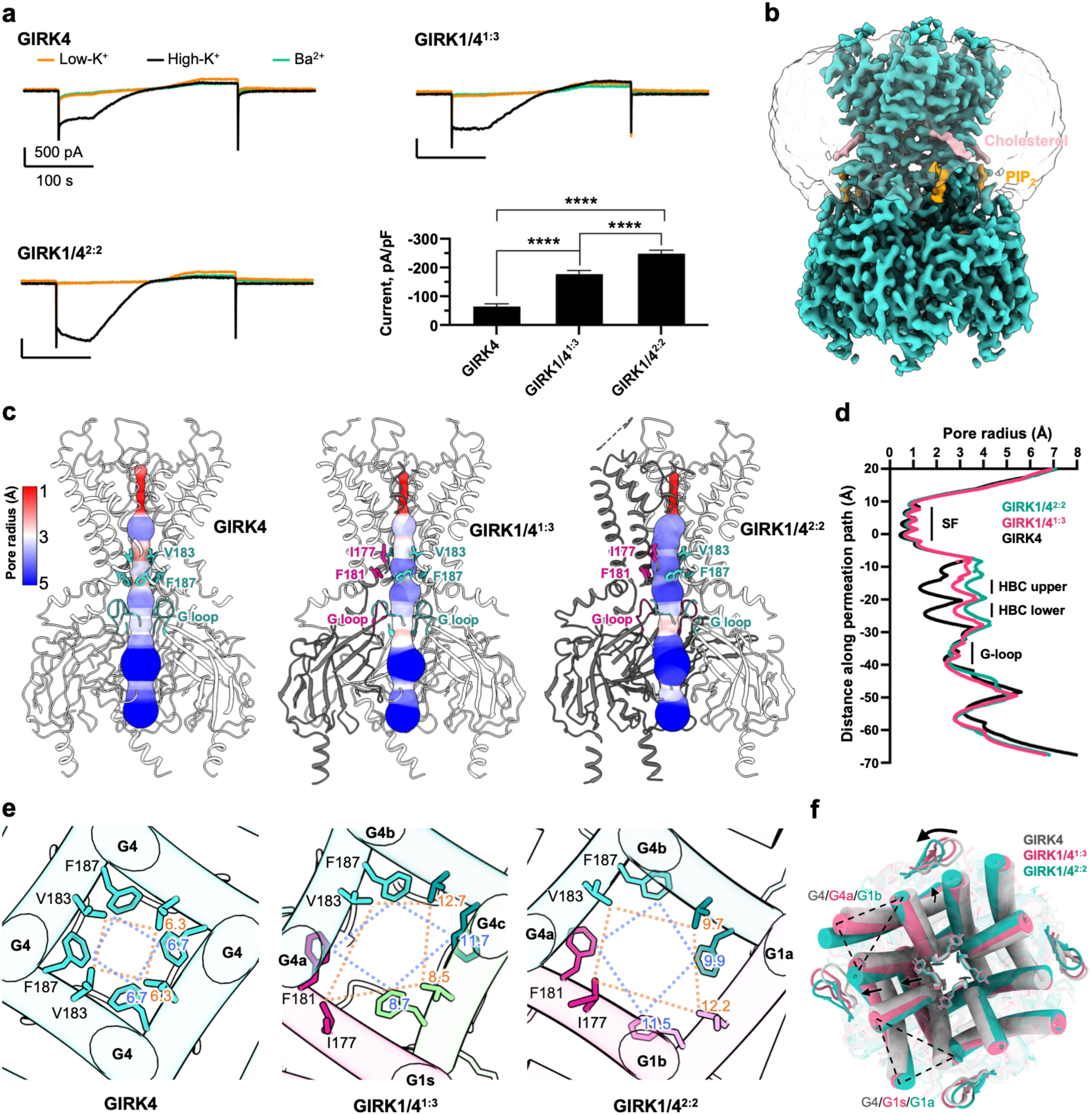
Distinct gating and pore architectures of GIRK4 and GIRK1/4 heterotetramers. **a**, Representative whole-cell patch-clamp current obtained in response to voltage-ramps from -80 to 80 mV in a perfusate containing 4 mM external K^+^ (low-K^+^, orange), 140 mM K^+^ (high-K^+^, black) and 140 mM K^+^ with 3 mM Ba^2+^ (Ba^2+^, green) for GIRK4 alone or the concatenated constructs indicated. The histogram shows the summary peak negative current-density in low and high K^+^ following subtraction of the Ba^2+^ inhibited current. Whole-cell patch-clamp was performed as described in the Methods. Summary data are from 6-10 cells per group from 3 independent experiments. **b**, Cryo-EM map of GIRK4 homotetramer. **c**, Pore radius profiles for GIRK4, GIRK1/4^1:3^ and GIRK1/4^2:2^ channels mapped onto the atomic models, highlighting constrictions at the HBC (upper and lower) and G-loop gates. **d**, Quantitative comparison of pore radius along the permeation pathway among GIRK4, GIRK1/4^1:3^, and GIRK1/4^2:2^, showing progressive gate opening with increasing GIRK1 stoichiometry. **e**, Top views of the HBC gate region. Yellow dashed lines connect residues forming the HBC upper gate, and blue dashed lines connect residues forming the HBC lower gate. The values indicate the shortest distances between opposing side chains that define the upper and lower constrictions. **f**, Superposition of GIRK4, GIRK1/4^1:3^, and GIRK1/4^2:2^ viewed from the extracellular side reveals a counterclockwise rotation of CTD, with the LM loops highlighted, as the GIRK1 content increases, corresponding to progressive opening of the HBC gate (see also fig. S8D).

To define the structural basis for the graded increase in basal activity, we determined the structure of the human GIRK4 homotetramer at 2.8 Å resolution (Fig. 2b and Extended Data Fig. 7) and compared its pore architecture with those of the two GIRK1/4 heterotetramers. Owing to symmetry, the GIRK4 homotetramer possesses a straight ion-conduction pathway, whereas the GIRK1/4 heterotetramers exhibit local curvature in the pathway due to the asymmetry introduced by GIRK1 (Fig. 2c). Quantitative comparison of pore dimensions reveals pronounced differences below the selectivity filter. In GIRK4, the HBC lower gate (formed by a phenylalanine) and the HBC upper gate (formed by an isoleucine in GIRK1 and a valine in GIRK4) both exhibit narrow constrictions that cannot accommodate hydrated K⁺ (Fig. 2d). Replacement of one GIRK4 subunit by GIRK1 in the GIRK1/4^1:3^ substantially increases the pore size at both HBC positions to approximately twice that of the homotetramer, whereas incorporation of a second GIRK1 subunits in GIRK1/4^2:2^ further enlarges the HBC gate (Fig. 2d,e).

GIRK1 incorporation also introduces pronounced local asymmetry at the HBC gate. In GIRK1/4^1:3^, the minimal distance is defined by two opposing GIRK4 side chains, whereas in GIRK1/4^2:2^ the GIRK1–GIRK4 pairwise distances are unequal, with G1b and G4b defining the minimum constriction (Fig. 2e). In contrast, the dimensions of the G-loop gate do not differ substantially among the three structures (Fig. 2d and Extended Data Fig. 8a-c), making it unlikely that the G-loop accounts for the graded increase in basal activity. Because the HBC gate is formed by bulky hydrophobic residues, changes in its dimensions are expected to alter the energetic barrier to hydrated K⁺ permeation. We therefore interpret the GIRK1-dependent increase in HBC size not as the sole determinant of basal activity, but as one structural component of broader inner-pore remodeling. Consistent with this interpretation, progressive GIRK1 incorporation drives outward splaying of the transmembrane helices near their cytoplasmic ends and is accompanied by a counterclockwise rotation of the CTD (Fig. 2f and Extended Data Fig. 8d).

All GIRK4 structures also exhibited a well-defined hydrogen-bond network surrounding Glu147, a residue critical for K^+^ permeation. Glu147 forms direct hydrogen bonds with Gln171 and Thr100, while Gln171 is further stabilized through H-bond interactions with Trp108 and Ser143. In contrast, GIRK1 lacks the equivalent hydrogen-bond network surrounding Glu141 due to two differences from GIRK4, Phe137 instead of Ser143 and Ala94 replacing Thr100. Similar differences, like in GIRK1/2 heteromers^67^, have been shown to contribute to the enhanced activity of GIRK1-containing channels relative to homomeric counterparts^32,33^ (Extended Data Fig. 9). Thus, incorporation of GIRK1 reshapes multiple structural features of the conduction pathway, combining progressive HBC widening with local remodeling of hydrogen-bond network surrounding the inner pore.

### Distinct modes of ML297 binding in GIRK1/4

ML297, engages GIRK1-centered intersubunit pockets whose local composition differs according to GIRK1/4 stoichiometry. The compound, a PIP₂-dependent selective agonist of GIRK1 heteromers^30,34^, contains an N-methylpyrazole–phenyl head group, a urea linker, and a difluorophenyl tail, forming a three-segment scaffold (Fig. 3a). In the ML297-bound reconstructions, the ligand engages the single GIRK1 subunit in the GIRK1/4^1:3^ (Fig. 3b), and either of the two GIRK1 subunits in the GIRK1/4^2:2^ complex (Fig. 3c,d). We did not capture a state with ML297 simultaneously bound to both GIRK1 subunits under our experimental conditions. In all structures, the binding pocket is primarily formed by the GIRK1, with additional contributions from the adjacent TM2 helix, which can originate from either GIRK4 or GIRK1 (Fig. 3b-d).

**Fig. 3.**
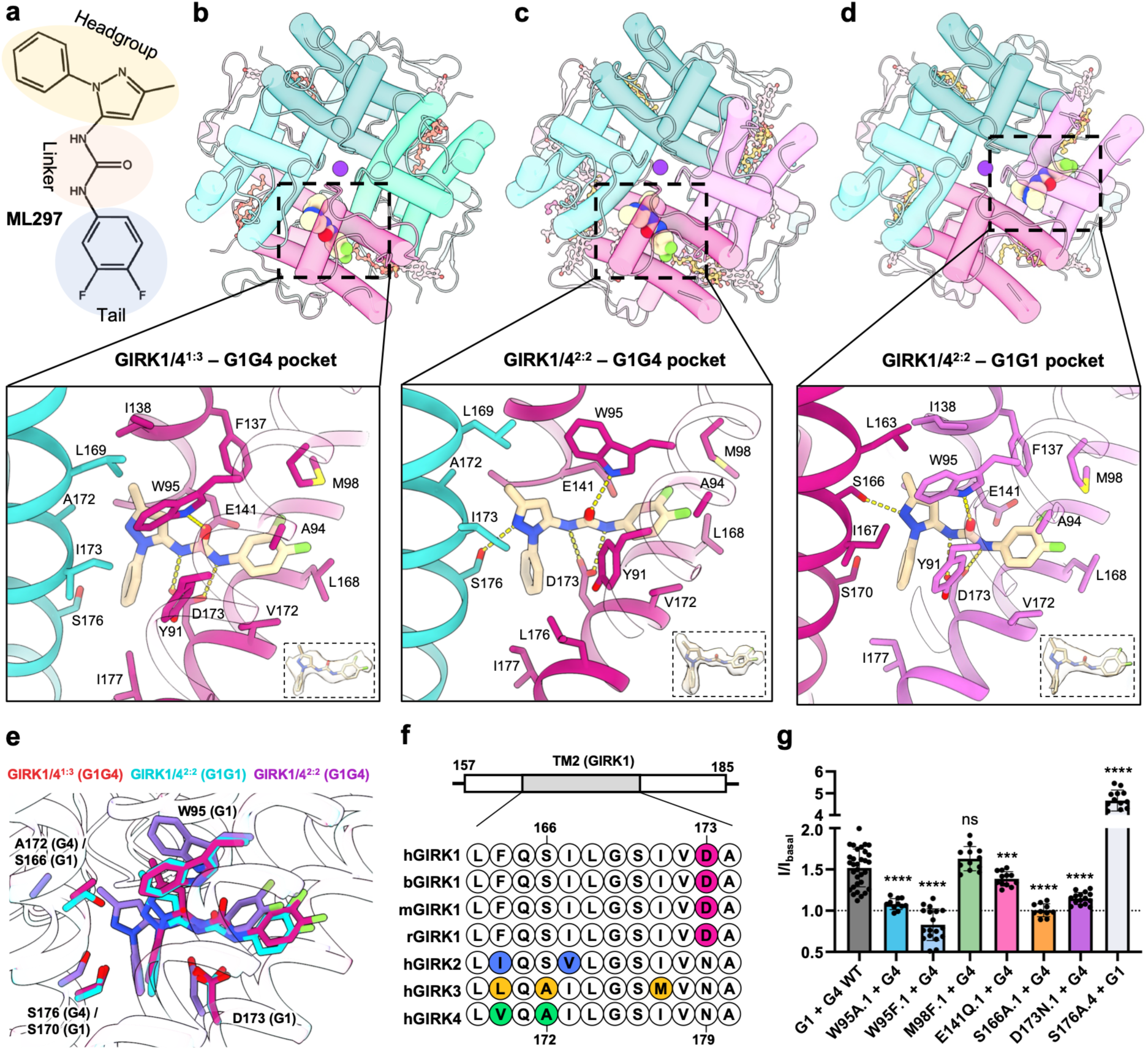
Structural basis of ML297 binding in GIRK1/4 heterotetramers. **a**, Chemical structure of the GIRK1/4 activator ML297. **b**-**d**, Top views and close-up views of the ML297 binding pockets in GIRK1/4^1:3^-ML297 (**b**), GIRK1/4^2:2^-ML297(G1G4) (**c**), GIRK1/4^2:2^-ML297(G1G1) (**d**). ML297 is coordinated through a network of hydrogen bonds and hydrophobic contacts involving residues from both subunits at the interfacial pockets. Insets show the well-defined density of ML297. **e**, Superposition of ML297-binding pockets from GIRK1/4^1:3^ and GIRK1/4^2:2^ structures, highlighting conserved pocket features and candidate hydrogen-bonding positions relative to ML297. **f**, Sequence alignment of the TM2 residues forming the ML297 pocket from human (h), bovine (b), mouse (m), and rat (r) GIRK1, together with human GIRK2–4. Non-conserved residues are color-coded: magenta (GIRK1), blue (GIRK2), yellow (GIRK3), and green (GIRK4). **g**, Functional validation of key pocket residues by mutagenesis using *Xenopus* oocytes as the expression system. Mutations are indicated using the format XnY.subunit, where n denotes the residue position and the suffix denotes the GIRK subunit (.1 for GIRK1 and .4 for GIRK4). Point mutations in the ML297 site reduce (D173N.1, W95A.1, W95F.1, S166A.1, E141Q.1), while one site increases (S176A.4) drug-induced activation and one site shows no effect (M98F.1) (mean ± s.e.m.).

In the ML297-bound GIRK1/4^1:3^ structure, the compound adopts an overall pose similar to that we report in the ML297-bound GIRK1/2^1:3^ complex^67^. GIRK1 Trp95 engages with the carbonyl oxygen and Asp173 with the two nitrogens of the urea linker, while Met98 lies in the proximity to the tail group and Glu141 lies near the urea linker. A key difference arises from the adjacent subunit: Ser177 of GIRK2 is replaced by Ala172 in GIRK4, thereby eliminating the hydrogen bond to the N-methylpyrazole head group nitrogen observed in GIRK1/2^67^. Consistent with the loss of this stabilizing interaction, ML297 exhibits less defined density than in GIRK1/2, suggesting greater ligand mobility within the GIRK1/4^1:3^ pocket (Fig. 3b).

ML297 adopts a distinct configuration in the G1G4 pocket of GIRK1/4^2:2^. Here, the ligand shifts downward and forms a stable hydrogen bond with Ser176 of the GIRK4 subunit, located approximately one helical turn below Ala172 (Fig. 3c). Multiple cholesterol molecules are also resolved along the membrane-facing surface of this tetramer, where they form a stacked assembly with the channel (Extended Data Fig. 10a). One cholesterol molecule inserts into the inter-subunit cleft between G4a and G1b and contacts ML297 as well as surrounding hydrophobic residues from both subunits (Extended Data Fig. 10b).

As this pocket is absent from the corresponding structure without ML297, ligand binding may promote formation of this site, with cholesterol further stabilizing ML297 engagement. These observations indicate that the G1G4 pocket in the 2:2 heterotetramer is conformationally plastic, enabling a more stably engaged ML297 state that is accompanied by interfacial cholesterol binding. The G1G1 pocket in GIRK1/4^2:2^ provides a third local binding environment. In this pocket, Ser166 of the adjacent GIRK1 subunit, equivalent to Ser177 in GIRK2, forms a hydrogen bond with ML297. However, unlike the G1G4 pocket in GIRK1/4^2:2^, the G1G1 pocket adopts a conformation that more closely resembles that observed in the GIRK1/4^1:3^ complex (Fig. 3b,d,e). In both ML297-bound GIRK1/4^2:2^ structures, GIRK1 Trp95 forms a hydrogen bond with the urea-linker oxygen of ML297, while surrounding hydrophobic residues stabilize the nonpolar portions of the ligand. Notably, Asp173 of GIRK1—a residue unique to GIRK1 that is poorly resolved in the ML297-bound GIRK1/2 structure^67^ due to its conformational flexibility — is well resolved in the ML297-bound GIRK1/4 and forms a hydrogen bond with the urea-linker nitrogen (Fig. 3f).

We next tested the functional importance of these contacts by mutating interacting residues either on the ML297-bearing GIRK1 subunit or on the adjacent TM2 helix that completes the intersubunit binding pocket. Mutant subunits were co-expressed with their wild-type partner and tested in *Xenopus* oocytes (Fig. 3g and Extended Data Fig. 11a,b) and HEK293 cells (Extended Data Fig. 11c). Mutations disrupting direct ligand contacts generally reduced ML297-evoked activation in both expression system, although the magnitude of the effects differed in some cases. W95A/F.1 produced the most pronounced loss of potentiation, consistent with its central role in anchoring the urea linker. M98F.1 had little to no effect, while E141Q.1 and D173N.1 produced more modest reductions, indicating secondary or context-dependent contributions from these residues. Strikingly, mutation of Ser166 on the adjacent GIRK1 subunit (S166A.1), which contributes TM2 to the G1G1 pocket, nearly abolished ML297-dependent activation in both oocytes and mammalian cells (Fig. 3g and Extended Data Fig. 11). Because Ser166 contacts ML297 specifically within the G1G1 pocket in the resolved structures, this result identifies engagement of the G1G1 interface as a major determinant of ML297-induced channel activation. By contrast, S176A.4 enhanced potentiation, suggesting that local side-chain constraints at the G1G4 interface modulate ligand efficacy. Per-residue interaction energy estimates independently identify the same contact network as important for ML297 action (Extended Data Fig. 12). Mutations that altered ML297 responses preserved surface expression suggesting the changes in responses reflected functional changes and were not due to trafficking defects (Extended Data Fig. 13).

Structural superposition further reveals pocket-dependent conformational responses to ML297. The GIRK1/4^1:3^-ML297 and GIRK1/4^2:2^-ML297(G1G1) complexes are globally similar and position the ligand in nearly identical orientations. By contrast, in GIRK1/4^2:2^-ML297(G1G4), the TMD undergoes a slight outward displacement relative to the CTD, producing an expanded pocket that accommodates the downward-shifted ligand (Fig. 3e and Extended Data Fig. 14a). This rearrangement is accompanied by a change in the rotamer conformation of the conserved GIRK1 Trp95, suggesting a local structural adaptation associated with ligand repositioning (Fig. 3c,e). Comparison of structures with and without the ligand further shows that upon ML297 binding to the G1G4 pocket is accompanied by a clockwise TMD twist of ∼4° and ∼3° in the 1:3 and 2:2 stoichiometries, respectively, whereas binding to the G1G1 pocket of GIRK1/4^2:2^, is associated with ∼2 Å TMD displacement (Extended Data Fig. 14b).

These pocket-specific rearrangements extend to the HBC gate. Pore radius analysis shows that the selectivity filter and the G-loop gate remain largely unaffected upon ML297 binding (Extended Data Fig. 14c). In contrast, both the upper (Val/Ile) and lower (Phe) HBC constrictions are influenced. The largest gate expansion occurs in GIRK1/4^2:2^ with ML297 bound in the G1G1 pocket, followed by GIRK1/4^2:2^ with ML297 at the G1G4 pocket, whereas GIRK1/4^1:3^ exhibits the least HBC gate opening (Extended Data Fig. 14c). Thus, the local composition of the ML297-binding interface is coupled to distinct structural responses at the channel gate.

### ML297 drives opposite functional responses in the two GIRK1/4 stoichiometries

We next asked whether the distinct structures and binding environments of the two GIRK1/4 stoichiometries result in different pharmacological responses to ML297. We tested concatenated constructs that generate defined GIRK1/4^1:3^ and GIRK1/4^2:2^ populations (Fig. 4a-d). Remarkably, ML297 produced opposite effects on the two channels. In GIRK1/4^1:3^, ML297 inhibited basal high-K^+^ currents by an average of 48% with an IC_50_ of 0.23 μM (Fig. 4a,c,d). In contrast, ML297 increase GIRK1/4^2:2^ currents by an average of 73% with an EC_50_ of 0.72 μM (Fig. 4b-d). Thus, GIRK1/4 stoichiometry determines whether ML297 acts as an inhibitor or activator.

**Fig. 4.**
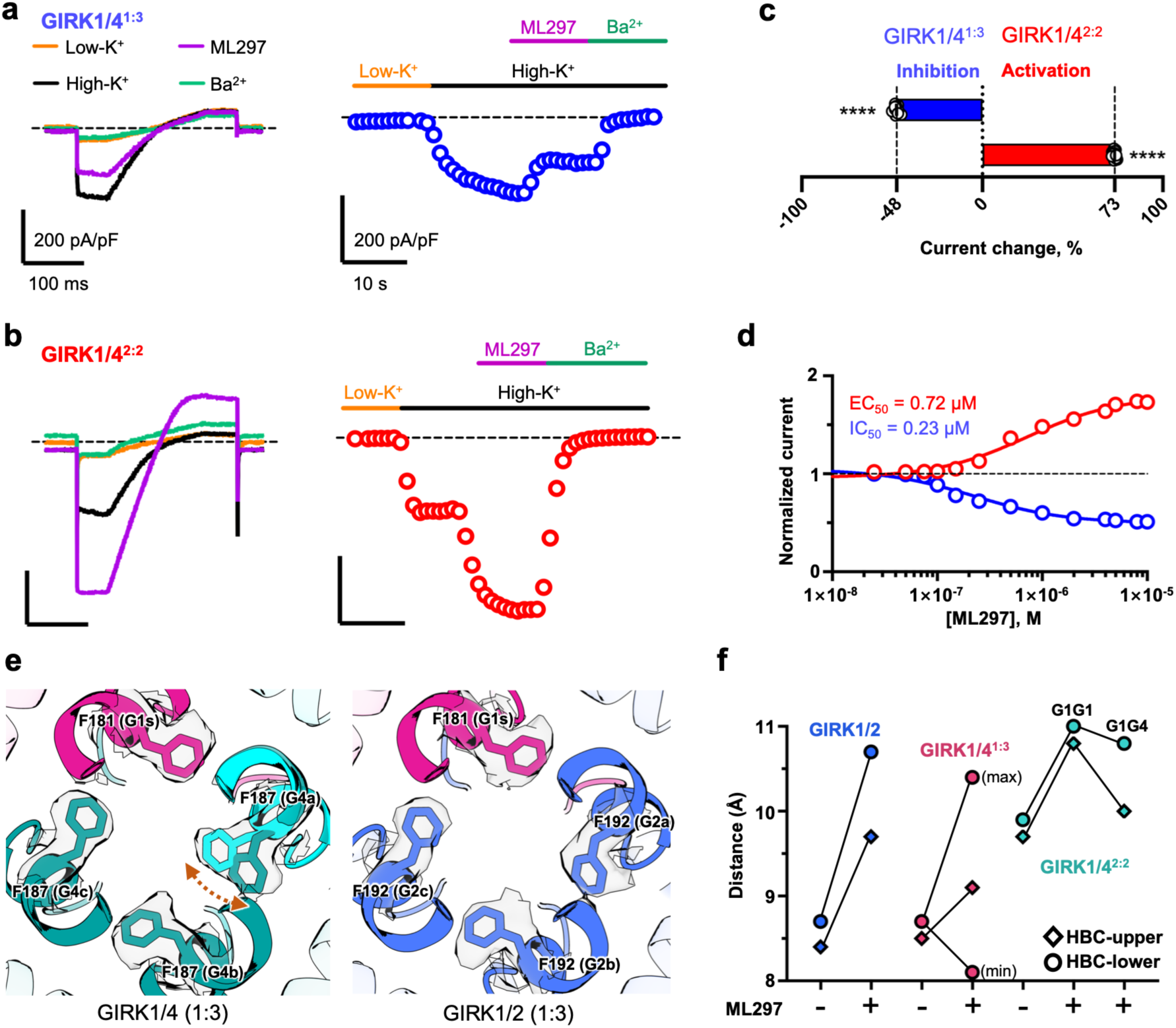
Stoichiometry-dependent effects of ML297 on GIRK1/4 channels. **a**,**b**, Representative whole-cell currents recorded from HEK293 cells expressing concatenated GIRK1/4 tetramers: GIRK1/4^1:3^ (**a**) or GIRK1/4^2:2^ (**b**) stoichiometries, showing opposite responses to ML297 application. Left, Representative current-sweeps; right, time-course plots (right) illustrate inhibition (blue) or activation (red) by 10 μM ML297. **c**, Summary of the change in current observed with each GIRK1/4 concatemer in response to 10 mM ML297. GIRK1/4^1:3^ current is inhibited by 48%, GIRK1/4^2:2^ current is increased by 73%. **d**, Summary concentration-response curves for GIRK1/4 concatemeric. ML297 inhibits GIRK1/4^1:3^ with an IC_50_ of 0.23 µM while GIRK1/4^2:2^ is increased with an EC_50_ of 0.72 µM. **e**, Structural comparison of HBC gates in GIRK1/4^1:3^ and GIRK1/2 channels viewed from the extracellular side, highlighting the large movements of Phe187 from G4a in GIRK1/4^1:3^ structure upon ML297 binding. The density maps are contoured at 5σ after normalization. **f**, Quantification of distances between opposing residues at the upper and lower HBC gates measured from structures with and without ML297. ML297 expands the HBC gate in GIRK1/2 and two GIRK1/4^2:2^ structures but could expand or narrow the HBC gate in GIRK1/4^1:3^ structure. Whole-cell patch-clamp was performed as described in the Methods. Summary data are from 6-10 cells per group from 3 independent experiments. Statistical significance was assessed using a two-tailed, unpaired Welches T-test where **P < 0.01 and ****P < 0.0001.

To explore the structural basis for inhibition of GIRK1/4^1:3^, we compared this complex with GIRK1/2^67^. Although both channels adopt a 1:3 stoichiometry, the side chain density of Phe187 in the GIRK4 subunit adjacent to GIRK1 (G4a) is substantially more diffuse than that of Phe192 on the corresponding GIRK2 subunit (G2a) of GIRK1/2^67^ (Fig. 4e). Together with the less well-defined ML297 density in GIRK1/4^1:3^ (Fig. 3b), this suggests that ML297 is less stably engaged in this interface. The instability reflects, at least in part, loss of the stabilizing hydrogen bond provided by the corresponding serine in GIRK2, because Ala172 in GIRK4, cannot anchor the N-methylpyrazole–phenyl head group. ML297 binding is therefore associated with continued local rearrangements at the G1s–G4a interface and pronounced wobbling of the Phe187 side chain in G4a (Fig. 4e).

We next quantified the minimal heavy-atom distances at the helix bundle crossing (HBC) gate in GIRK1/2 and in both GIRK1/4 stoichiometries (Fig. 4f). In the presence of PIP_2_, ML297 binding markedly widens both the upper and lower HBC gates in the GIRK1/2–ML297 and GIRK1/4^2:2^– ML297 (G1G1) structures, with the latter displaying the largest HBC opening observed (∼11 Å). In contrast, the GIRK1/4^1:3^ complex exhibits the smallest opening at the upper HBC gate. At the lower gate, however, swinging of Phe187 from the G4a subunit produces a heterogeneous opening, with maximal and minimal distances of 10.4 Å and 8.1 Å, respectively. This uncoupled behavior of the upper and lower HBC constrictions, which alternate between widened and restricted conformations relative to the activator-free state, provides a structural correlate of the net inhibitory effect of ML297 on GIRK1/4^1:3^.

More generally, the structures indicate that ML297 engagement of a G1G1 interface most strongly promotes HBC opening, whereas engagement of a G1G4 interface produces a smaller and more variable effect. This distinction provides a structural explanation for both the opposing responses of the two stoichiometries and previous observations that ML297 activates GIRK1/4 less efficiently than GIRK1/2, with a lower E_max_ and a higher EC_50_ for GIRK1/4^34^.

Gating molecular dynamics (GMD) simulations further support this pocket-dependent pharmacology. Aggregate simulations totaling 3 µs were performed for all three ML297-bound GIRK1/4 structures (Extended Data Fig. 15). Across all complexes, the ML297 urea linker maintained persistent interactions with GIRK1 Asp173 (D173.1), with this contact present for 63.5% of the simulation time in the GIRK1/4^1:3^-G1G4 pocket, 95% in the GIRK1/4^2:2^-G1G4 pocket, and 77% in the GIRK1/4^2:2^-G1G1 pocket. In contrast, ML297 showed minimal engagement with GIRK1 Glu141 (E141.1) in all GIRK1/4 simulations, differing from its behavior in ML297-bound GIRK1/2^67^.

The contribution of GIRK1 Trp95 (W95.1) to ligand coordination was particularly dependent on the pocket identity. In the 1:3 G1G4 pocket, W95.1 contacted the urea linker for only 26.9% of the simulation time, whereas in both 2:2 pockets this interaction persisted for more than 70% of the simulations, approaching the persistence of the D173.1 interaction. These differences correlated with stabilization of the ML297 pyrazole head group. In the 2:2 G1G1 pocket, Ser166.1 formed a direct hydrogen bond with the pyrazole nitrogen, whereas in the 2:2 G1G4 pocket a downward-shifted ligand retained a hydrogen bond with Ser176.4. Cholesterol appeared to stabilize interactions involving W95.1 and D173.1, while weakening the S176.4 in the 2:2 G1G4 pocket. By contrast, the ligand head group formed only transient contacts with S176.4 in the 1:3 G1G4 pocket, and this reduced head-group stabilization was accompanied by weaker W95.1 engagement and lower ion permeation. Together, the simulations support the structural and electrophysiological evidence that differences in intersubunit pocket composition alter ML297 pose stability and its coupling to channel conduction.

### Extensive water–protein interactions in GIRK1/4

The high-resolution cryo-EM map of the GIRK1/4^1:3^ complex enables direct visualization of ordered water molecules and bound ions, providing a detailed view of the hydration network surrounding key functional elements of the channel (Fig. 5). Structured waters form hydrogen-bonding networks that connect spatially separated regions of the protein and may contribute to their conformational stability^35–38^. To ensure confident assignments, water sites were assigned using stringent density- and geometry-based criteria in *phenix.douse* and validated by Q-score analysis (mean = 0.89; minimum > 0.75), indicating high local resolvability (Fig. 5b). Independent evaluation using the deep-learning–based MIC^39^ classifier further supported these assignments (Fig. 5c).

**Fig. 5.**
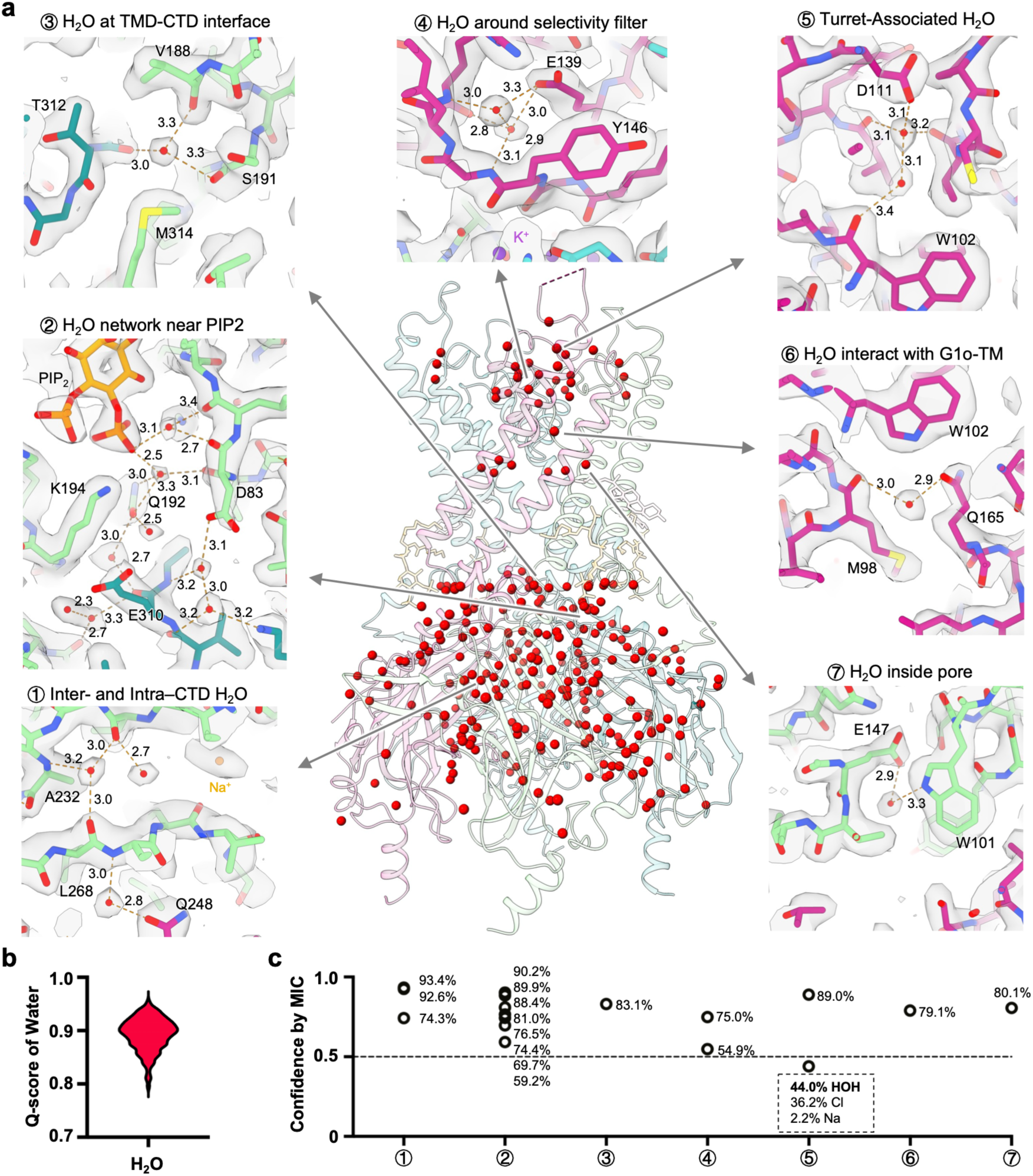
Hydration network in the GIRK1/4 heterotetramer. **a**, Central panel, overall distribution of water molecules (red spheres) modeled into the GIRK1/4^1:3^ structure. Arrows indicate representative hydration site shown in panels 1-7; distances (Å) from water to protein heavy atoms are labelled. 1, Inter- and intra-CTD water molecules mediating backbone interactions. 2, Hydration network near PIP_2_ involving basic residues and phosphates. 3, Water cluster at the TMD-CTD interface forming hydrogen-bond networks with polar side chains. 4, Water molecules coordinated around the selectivity filter. 5, Turret-associated water stabilized by acidic and aromatic residues. 6, Water bridging the G1o transmembrane helices. 7, Water molecules located inside the ion-conduction pore. **b**, Distribution of water Q-scores. **c**, Confidence values of individual waters predicted by the MIC program, indicating the probability of each site being assigned as H_2_O (HOH) or ions.

Most ordered waters localize to the CTD, where they mediate extensive intra- and inter-subunit hydrogen-bonding interactions that may stabilize the CTD fold and quaternary organization (Fig. 5a). Additional water-binding sites cluster around the Na⁺-binding pocket, suggesting a contribution of local hydration to the electrostatic environment that underlying Na⁺-dependent regulation of GIRK activity.

Structured Waters are also positioned at interfaces that couple the TMD and CTD. Whereas PIP₂ is known to bridge these domains directly^22,31^, the GIRK1/4 structure reveals additional water-mediated PIP_2_–protein interactions. At the TMD–CTD interface, structured waters coordinate G-loop residues (Ala311 in GIRK4; Thr305 in GIRK1) with TM2 residues near the cytoplasmic end (Val188 and Ser191 in GIRK4; Ile182 and Ser185 in GIRK1), potentially reinforcing the docked state. Near the selectivity filter, two conserved waters per subunit bridge a negatively charged glutamate (Glu139 in GIRK1; Glu145 in GIRK4) to the backbone of the pore-loop backbone. Together with turret-associated waters, these interactions may buffer local electrostatics and mechanically stabilize the selectivity filter and extracellular loop region during gating transitions.

Ordered waters are also resolved within the transmembrane region. A surface-exposed water molecule in GIRK1 bridges TM1 and TM2, while a water molecule in the central cavity hydrogen-bonds with Glu147 and Trp101 in GIRK4, positioning it to participate in K⁺ hydration and dehydration during permeation. Collectively, these observations define an interconnected hydration network linking lipid binding, gating, and ion-selective elements and support a role for structured water in GIRK channel conduction and regulation.

## Discussion

Our findings establish subunit stoichiometry as an important determinant of both gating and ligand responsiveness in cardiac GIRK channels. Rather than forming a single canonical heterotetramer, GIRK1 and GIRK4 assemble into a compositional ensemble that includes asymmetric 1:3 and parallel 2:2 GIRK1/4 heterotetramers together with symmetric GIRK4 homotetramers, The relative abundance of the heteromeric assemblies depends on subunit availability, with the 1:3 configuration predominating under the conditions examined here. Functionally, progressive GIRK1 incorporation is accompanied by increasing basal current and widening of the HBC gate without a comparable change in the G-loop gate. These observations provide a structural basis for the rank order of basal activity, GIRK4 < GIRK1/4^1:3^ < GIRK1/4^2:2^, and show how heteromer stoichiometry can tune the intrinsic gating properties of cardiac GIRK channels.

The ML297-bound structures further reveal that subunit stoichiometry alters pharmacology by changing the local intersubunit environment presented to the ligand. ML297 occupies a GIRK1-centered pocket that is completed by TM2 of the adjacent subunit, producing chemically and structurally distinct G1G1 and G1G4 interfaces. Binding within the 2:2 G1G1 pocket is associated with the largest HBC expansion and robust channel activation, consistent with mutagenesis identifying S166.1 of the adjacent GIRK1 subunit as an important determinant of ML297 efficacy. In contrast, G1G4 interfaces provide weaker or alternative head group stabilization, and the 1:3 complex exhibits net inhibition, associated with reduced W95.1 engagement and greater conformational heterogeneity near the lower HBC gate. GMD simulations further support a mechanism dominated by persistent D173.1 coordination, pocket-dependent W95.1 interactions and differential stabilization of the pyrazole head group. Thus, the same ligand can produce opposite functional responses depending on the subunit composition of the pocket through which it acts.

The GIRK1/4 structures also reveal additional lipid-associated interactions that may influence ligand engagement and channel gating. Cholesterol is known to modulate GIRK channels^40,41^ function, both directly through direct binding and through effects on membrane properties^42–44^. Beyond cholesterol sites identified previously in GIRK2 homotetramers^22^, the GIRK1/4^2:2^ structure containing ML297 in the G1G4 pocket reveals additional cholesterol molecules that directly contact and stabilize the channel-ligand complex. We also observed density near the PIP₂-binding site, that can be fitted with the maltoside headgroup of GDN molecule (Extended Data Fig. 16a). The headgroup forms hydrogen bonds with neighboring residues and the adjacent PIP_2_ molecule at the TM1 interface. A similar density is observed at this location across GIRK structures from this and previous studies (Extended Data Fig. 16b), raising the possibility that endogenous polar lipid headgroups may occupy an equivalent position in native membranes and reinforce PIP_2_ association. Although the physiological identity of such a lipid remains unresolved, this region defines an additional structural site through which PIP_2_-dependent gating could potentially be modulated.

The high-resolution GIRK1/4^1:3^ map further provides a detailed view of the electrostatic and hydration landscape through which ions enter and permeate the channel, ordered waters bridge PIP_2_ headgroups, coordinate Na⁺ ions, and connect key residues surrounding the selectivity filter and the TMD–CTD interface. These hydration networks may reinforce the PIP_2_-dependent docked state and stabilize elements of the conduction pathway. The explicit structural assignment alsoprovides an experimentally anchored framework for future molecular dynamics simulations aimed at defining how hydration contributes to ion entry, selectivity, permeation, and PIP_2_-dependent gating.

Together, these findings establish a stoichiometry-encoded mechanism of GIRK pharmacology in which subunit composition controls both gating energetics and the chemical environment of an allosteric ligand-binding pocket. Distinct GIRK1/4 assemblies can therefore respond differently, and even oppositely, to the same small molecule. This link between heteromer composition, pocket architecture, and ligand efficacy provides a structural framework for developing GIRK modulators that distinguish among channel assemblies and may ultimately enable greater selectivity between cardiac and neuronal GIRK channels.

## Methods

### Molecular Biology

*Biochemistry* – GIRK1/4 (hGIRK4-TEV-FLAG-p2A-hGIRK1-TEV-GFP-His10) and GIRK4 (hGIRK4-TEV-FLAG) in pEG_BacMam^45^ were generated via GenScript (Piscataway, NJ). BacMam baculoviruses were generated by the Bac-to-Bac System (Thermo Fisher Scientific, Waltham, MA).

*Electrophysiology* – The pMAX(+) (in-house vector) and pXoom^46^ vectors were used for expression in HEK293T cells and *Xenopus* oocytes. The pGEMHE^47^ vector was also used for *Xenopus* oocyte expression. Wild type and mutants of GIRK1 and GIRK4 were inserted in the pXoom or pGEMHE vectors. Mutations were introduced by QuikChange site-directed mutagenesis (Agilent, Santa Clara, CA) and confirmed by sequencing. GIRK1/4 concatemers in 1:3 and 2:2 stoichiometry (GIRK4-10Q-GIRK1-10Q-GIRK4-10Q-GIRK4 and GIRK4-10Q-GIRK1-10Q-GIRK1-10Q-GIRK4) in pMAX(+) were generated via GenScript (Piscataway, NJ). For TEVC using *Xenopus* oocytes, mRNAs were transcribed using mMESSAGE mMACHINE™ T7 Transcription Kit (Thermo Fisher Scientific, Waltham, MA) following plasmid linearization. *TIRF microscopy* – GIRK1-HA-mTFP1 and GIRK4-FLAG-TagRFP-T in pMAX(+) were ordered to GenScript (Piscataway, NJ).

### Protein Expression

Human GIRK4 and GIRK1/4 channels were expressed in HEK293S GnTI^-^ cells using baculovirus generated in-house. Suspension cultures of HEK293S GnTI^-^ cells were maintained at a density of 0.2 × 10^6^ cells/mL in a 37 °C incubator at 140 rpm with 8% CO_2_. When cultures reached a density of 2-3.5 × 10^6^ cells/mL, P4 baculovirus encoding the channel of interest was added and incubated for overnight. On the next day, sodium butyrate was added to a final concentration of 10 mM, and the incubation temperature was reduced to 30°C while maintaining 8% CO_2_. Cultures were incubated for an additional 48 hrs. Samples were collected daily from the time of transduction to monitor protein expression by Western blot. The cells were then harvested by centrifugation at 8,000 g for 10 min at 4°C.

### Membrane Preparation

Induced cell pellets were resuspended in ice-cold lysis buffer containing 50 mM Tris-HCL (pH 8), 200 mM NaCl, 20 mM KCl, 1 mM EDTA, 10% (v/v) glycerol and 1 mM PMSF, supplemented with a fresh protease inhibitor cocktail (Thermo Scientific, Cat. No. A32963). Cells were lysed with a microfluidizer at 4000 psi at 4°C. Cell lysate was centrifuged at 35,000 rpm for 30 min in a 45 Ti rotor (Beckman Coulter) to remove soluble protein fractions. The resultant pellet was resuspended in wash buffer containing 50 mM Tris-HCL (pH8), 500 mM KCl, 1 mM EDTA, 10% glycerol, 1 mM PMSF, and centrifuged again at 35,000 rpm for 30 min in a 45 Ti rotor (Beckman Coulter). The final membrane pellets were resuspended in solubilization buffer containing 50 mM Tris-HCL (pH 8), 150 mM KCl, 0.1 mM PMSF, supplemented with fresh protease inhibitors. Membrane protein concentration was determined using a BCA assay (Thermo Scientific, Cat. No. A55864). Membranes were either flash-frozen in liquid nitrogen and stored at -80°C or immediately used for detergent extraction.

### Detergent Extraction of Mammalian Cell Membranes and Purification

Membranes were diluted to 5 mg/mL in solubilization buffer and extracted with 1% (w/v) n-dodecyl-β-D-maltoside (DDM, Gold Biotechnology, Cat. No. DDM10) for 2 hrs at 4°C. The mixture was centrifuged at 18,600 rpm for 20 min in a 45 Ti rotor (Beckman Coulter). The supernatant containing solubilized membrane proteins was collected and incubated with pre-equilibrated Anti-DYKDDDDK G1 Affinity Resin (GenScript, Cat. No. L00432) for 2 hrs at 4°C. The resin was washed with 10 column volumes (CV) of wash buffer containing 20 mM Tris-HCl (pH 8), 150 mM KCl, 200 mM NaCl, 0.1% DDM, 20% glycerol, 0.1 mM PMSF, and 0.5 mM EDTA. Bound protein was eluted using the same buffer supplemented with 200 ug/mL DYKDDDDK-peptide (APExBIO, Cat. No. A6002). For GIRK1/4, an additional affinity-purification step using Ni-NTA agarose beads (GoldBio, Cat. No. H-320) was performed to select GIRK1-containing channels prior to purification using Anti-DYKDDDDK. Protein yield and purity were assessed by SDS-PAGE. High quality fractions were pooled and further purified by Size Exclusion Chromatography (SEC) using Superose6 Increase 10/300 GL (Cytiva, Cat. No. 29091596) column pre-equilibrated with 20 mM Tris-HCL (pH 8), 100 mM KCL, 0.01 ∼ 0.02% DDM, and 10 mM DTT, 1 mM EDTA. Peak fractions were concentrated using a 100 kDa MWCO concentrator (Millipore, Cat. No. UFC910008) to a ∼5 mg/mL and sent for cryo-EM analysis.

### Cryo-EM data collection and processing

Sample were evaluated by negative-stain EM^48^, prior to cryo-EM grid preparation. Cryo-EM grids were prepared using a Vitrobot Mark IV (Thermo Fisher Scientific). A volume of 3-4 µL of purified sample was applied onto glow-discharged 300-mesh R1.2/R1.3 UltrAuFoli holey gold grids (Quantifoil) under 100% humidity at 8 ℃. After 4 s of blotting, the grids were plunge-frozen in liquid ethane cooled by liquid nitrogen and then stored in liquid nitrogen.

Cryo-EM data were collected on a Titan Krios G3i (Thermo Fisher Scientific) transmission electron microscope equipped with an energy filter and a Gatan K3 direct electron detector. Movie micrographs were recorded using EPU software. For the GIRK1/4 sample, movies were collected with 50 frames, a total dose of 50 e^-^/Å^2^, a pixel size of 0.6485 Å, and defocus range of -0.8 to -1.8 µm. For the GIRK4 sample, movies were collected under similar conditions but with a pixel size of 0.92 Å.

For the GIRK1/4 without ML297 dataset, a total of 13,378 movie micrographs were collected. After patch motion correction and patch CTF estimation, template-based particle picking and extraction were performed for 2D classification, after multi-rounds of 2D classification, part of the selected particles was used to generate initial 3D references, then using all particles to do the 3D heterogenous refinement with the generated references in CryoSPARC^49^. The best 3D class was further refined by nonuniform refinement in CryoSPARC using the GIRK1/2 map as a reference to align the particles to their correct relative positions in the reconstruction. Then, further 3D classification was carried out in RELION^50^ to separate different stoichiometries. To improve map quality, the data were re-imported into CryoSPARC for nonuniform refinement and local refinement to achieve fast and accurate particle alignment. The refined particles were then re-exported to RELION for further 3D classification. Finally, the data were returned to CryoSPARC for final map reconstruction through CTF refinement, reference-based motion correction and map sharpening. A 2.48 Å map reconstructed from 44,397 particles with a 2:2 (GIRK1:GIRK4) assembly and a 2.28 Å map reconstructed from 91,528 particles with a 1:3 (GIRK1:GIRK4) assembly were obtained. A detailed workflow of data processing is provided in Extended Data Fig. 2.

For the GIRK1/4-ML297 dataset, a total of 13,007 movie micrographs were collected using the same strategy as described above. Three reconstructions were obtained: a 2.42 Å map from 92,036 particles with a 2:2 (GIRK1:GIRK4) assembly showing the ML297 density in the G1G4 pocket; a 2.77 Å map from 22,681 particles with a 2:2 (GIRK1:GIRK4) assembly showing ML297 density in the G1G1 pocket; and a 2.82 Å map from 25,050 particles with a 1:3 (GIRK1:GIRK4) assembly showing ML297 density in the G1G4 pocket.

For the GIRK4 dataset, a total of 3,445 movie micrographs were collected. After patch motion correction and patch CTF estimation, template-based particles picking and extraction were performed to generate particle images for 2D classification. Multiple rounds of 2D classification were followed by multiple rounds of heterogenous refinement and nonuniform refinement. The remaining particles were then imported into RELION for further 3D classification. The best class was re-imported into CryoSPARC for final map reconstruction through CTF refinement, reference-based motion correction and map sharpening. A 2.80 Å map reconstructed from 7,971 particles with C4 symmetry was obtained.

All the resolutions shown above are estimated by CryoSPARC with a cutoff for Fourier shell correlation (FSC) of 0.143.

### Model building and refinement

Initial structural models of GIRK1 and GIRK4 subunit predicted by AlphaFold^51^ were retrieved from the UniProt database. The models were first fitted into the final cryo-EM density map using UCSF Chimera, followed by manual adjustment in Coot^52^ and subsequent real-space refinement in Phenix^53^. The model quality was assessed in Molprobity^54^, and the refinement statistics are summarized in Extended Data Table 1.

For water modeling, the highest-resolution density map of GIRK1/4^1:3^ structure was processed using phenix.douse. To ensure confident water placement, a CCmask (cc_mask_filter) with the cutoff value of 0.7, a map threshold scale factor of 0.6, and a sphericity criterion (scc) of 0.7 were applied.

### Pore analysis and visualization

Pore volumes were created by MOLEonline^55^ and ChimeraX^56,57^, Pore radius along the permeation pathway was calculated by HOLE^58^. UCSF Chimera and ChimeraX were used for all structures visualization and figures preparation.

### Cell culture

Human embryonic kidney (HEK293T) cells were acquired from American Type Culture Collection (ATCC) and were maintained in Dulbecco’s modified Eagle’s medium (ATCC) supplemented with 100 units/ml penicillin, 100 μg/mL streptomycin, and 10% (vol/vol) fetal bovine serum. All cells were incubated in a 37 °C, humified incubator supplemented with 5% CO2. HEK293T cells were transfected using polyethylenimine solution (1 mg/ml) at a ratio of 6 µl per 1 µg of plasmid DNA, 24-30 hours prior to experiments. Cells for study were identified by expression of a fluorescence marker or via the encoded fluorescence tag indicated, typically eGFP for patch-clamp studies.

### Total internal reflection fluorescence (TIRF) microscopy

GIRK1 and GIRK4 were tagged at the C-terminus with mTFP1 and Tag-RFP-T, respectively, and imaged at the surface of live *Xenopus* oocytes cells by TIRF microscopy according to our prior methods^59,60^. Cells were studied after 24-32 hours in Low-K^+^ recording solution bath solution comprising, in mM: KCl 2, NaCl 96, MgCl_2_ 1, and HEPES 5, buffered to pH 7.4 with KOH. mTFP1 was excited by a 445-nm laser and Tag-RFP-T by a 561-nm laser (Coherent, Santa Clara, CA). The beams were conditioned for coherence with custom built Keplerian beam expanders upstream of laser cleanup filters (445/10 nm and 561/10 nm, respectively). Laser lines were tuned to provide 10 mW of incident light on a micro mirror positioned below a high numerical-aperture apochromat objective (60x, 1.5 NA; Olympus, Waltham, MA) mounted on an RM21 microscope frame equipped with a piezo-driven nanopositioning stage (MadCity Labs., Madison, WI). The emission of mTFP1 and Tag-RFP-T were isolated from the excitation beam by an exit micro mirror and a ring-diaphragm positioned below the micro mirror assembly. When the fluors were imaged simultaneously, the emission was split by 510-nm dichroic mirrors placed downstream of parallel 480/40 nm and 620/60 nm bandpass filters. The fluorophores were imaged using a back illuminated sCMOS camera (Teledyne Photometrics, Tucson, AZ) controlled by µ-Manager software^61^. Additional lenses were placed downstream of the camera to adjust the final magnification of the emission image to 100 nm / pixel. All filters and mirrors were from Chroma (Bellow Falls, VT). Lenses, pinholes and diaphragms were from ThorLabs (Newton, NJ). Tetrapsec beads (Thermo, Waltham, MA) were routinely imaged to map the sCMOS chip and to calibrate the evanescent field depth to 100 nm. To assess stoichiometry, fluorophores were photobleached by continual excitation. Data were captured as 16-bit movies of up to 600 frames acquired at 1 Hz. Emission from each fluorophore was captured simultaneously to a mapped sCMOS chip, saved as separate image stacks and processed in an identical manner. Images were corrected for background fluorescence by subtracting the mean of 5 fully bleached frames from the end of each stack analyzed^62^. Fluorescent particles were defined as a discrete 3 x 3-pixel region around a pixel of maximum intensity, as before, and misalignment was corrected in ImageJ using StackReg. Co-localization of partner pixels from the two stacks of images was defined as the presence of both fluorophores with at least 30% of maximum fluorescence levels recorded in that region of interest, as before^60^.

### Single-particle photobleaching analysis of GIRK1/4 stoichiometry

Experimental design: Single-particle TIRF photobleaching was performed in 3-5 independent experiments using separate batches of Xenopus oocytes on separate days, each comprising at least three biological replicates. (1) Co-injection experiments: oocytes were injected with both GIRK1-mTFP1 and GIRK4-TagRFP-T cRNA at the indicated ratios. Particles colocalized between the two fluorescence channels (k₁ ≥ 1 and k₂ ≥ 1) were scored for heteromer stoichiometry analysis. (2) GIRK4-alone experiments: oocytes were injected with GIRK4-TagRFP-T cRNA alone, with water substituted for GIRK1 cRNA. TagRFP-T-positive membrane particles (k₁ = 0, k₂ ≥ 1) were scored as a positive control for tetrameric stoichiometry. (3) GIRK1-alone experiments: oocytes were injected with GIRK1-mTFP1 cRNA alone, with water substituted for GIRK4 cRNA. mTFP1-positive membrane particles (k₁ ≥ 1, k₂ = 0) were scored as a qualitative test of GIRK1 trafficking in the absence of an obligate partner subunit.

Particle classification and scoring: In all three experiments, fluorescence-versus-time records for individual particles were scored by manual review as having k distinct, monotonic bleaching steps above background, with a check that successive steps were of similar magnitude. Particles were tabulated by their joint (k₁, k₂) labels for the co-injection experiments and by their single-channel (k₂) or (k₁) labels for the GIRK4-alone and GIRK1-alone experiments respectively.

Zero-truncated binomial framework: Under the standard photobleaching framework^63^, a tetrameric channel containing n fluorescently labelled subunits with per-fluorophore step-detection probability p produces a binomial distribution of observed step counts. Because particles can only be detected if at least one step is observed (k ≥ 1), the relevant distribution is the zero-truncated binomial:

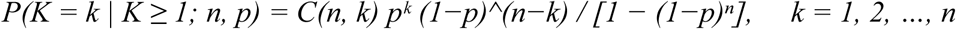

where C(n, k) = n! / [k!(n−k)!] is the binomial coefficient. The numerator gives the standard binomial probability of observing k steps from n labelled subunits with detection probability p; the denominator removes the probability mass of the unobserved zero-step outcome and renormalizes the distribution. The detection probability p reflects the probability that a single bleaching event is correctly resolved as a discrete step against background noise, and combines the underlying fluorophore quantum yield with collection efficiency, integration time, and the criteria used to score steps; p is fit from the data rather than assumed.

Heteromer mixture model: For the co-injection experiments, heteromer step distributions were fit to a bivariate zero-truncated binomial mixture comprising three components representing the three possible heteromeric stoichiometries: 1 GIRK1 : 3 GIRK4 (n₁ = 1, n₂ = 3), 2:2 (n₁ = 2, n₂ = 2), and 3:1 (n₁ = 3, n₂ = 1). Under this model, the probability of observing (k₁, k₂) bleaching steps in a localized particle of stoichiometry (n₁, n₂) is

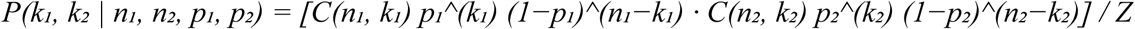

with normalisation Z = [1 − (1−p₁)^(n₁)] [1 − (1−p₂)^(n₂)] reflecting the colocalization gate criteria, i.e. k₁ ≥ 1 and k₂ ≥ 1. The full predicted joint distribution per condition is

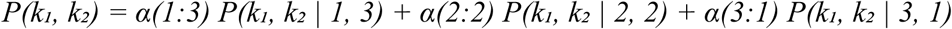

with mixing weights α ≥ 0 and α(1:3) + α(2:2) + α(3:1) = 1. The detection probabilities p₁ (for GIRK1-mTFP1) and p₂ (for GIRK4-TagRFP-T) were treated as shared across all conditions; mixing weights were free per condition. The model was fit by maximum likelihood across the five cRNA conditions in which heteromer assembly proceeds normally (1:1, 1:2, 2:1, 1:9, 9:1) using scripts in R established by Hines, 2013^63^. cRNA mixing ratios (1:49 and 49:1) were excluded from the formal fit because reduced membrane particle counts and skewed assembly under these conditions limit the inference. 95% confidence intervals on all parameters were obtained by parametric bootstrap (100 replicates) resampled from the fitted mixture. Goodness-of-fit was assessed by Pearson χ² on the six observed (k₁, k₂) bins per condition, with bin merging where expected counts fell below 5. The analysis framework follows Hines (2013), extended to a bivariate mixture, and is consistent with our recent application of the framework to Kir2.1– SUMO1 stoichiometry^64^.

GIRK4-alone positive control: As a positive control on a known-tetrameric population, GIRK4-TagRFP-T was expressed in Xenopus oocytes by injection of GIRK4 cRNA alone, with water substituted for GIRK1 cRNA. The experiment was performed on separate days and using separate biological batches of oocytes from the co-injection experiments. TagRFP-T-positive membrane particles (k₁ = 0, k₂ ≥ 1) were scored as above. Step counts were pooled across all cRNA-dose batches and fit to a zero-truncated binomial with n = 4 by maximum likelihood, with 95% bootstrap CIs and Pearson χ² goodness-of-fit on the four observed bins. This experiment validates that GIRK4 alone traffics to the plasma membrane as a tetramer and that the photobleaching framework correctly recovers a 4-subunit binomial distribution on a known-tetrameric population.

GIRK1-alone trafficking observation: As a control for the trafficking properties of GIRK1, GIRK1-mTFP1 cRNA was injected into Xenopus oocytes alone, with water substituted for GIRK4 cRNA, on separate days and using separate biological batches from the co-injection and GIRK4-alone experiments. mTFP1-positive membrane particles (k₁ ≥ 1, k₂ = 0) were tabulated. No quantitative binomial fit is reported for this dataset given the small particle count; the experiment is reported as a qualitative observation that GIRK1 traffics poorly to the plasma membrane in the absence of an obligate partner subunit.

### Two-electrode voltage-clamp

The plasmid DNA for each channel construct was linearized prior to *in vitro* transcription and transcribed using an mMessage mMachine T7 transcription kit (Thermo Fisher Scientific). *Xenopus* oocytes were surgically extracted in accord with an IACUC protocol at Northeastern University. Oocytes were then dissociated and defolliculated by collagenase treatment and microinjected with 50 nL of an RNAse/ DNAse free water solution per oocyte containing 1ng of cRNA. Injected oocytes were incubated for 1-2 days at 17 °C before currents were recorded. Electrodes were pulled using a Flaming-Brown micropipette puller (Sutter Instruments) and were filled with 3 M KCl in 1.5% (w/v) agarose to give resistances between 0.5 and 1.0 MΩ. The oocytes were bathed in ND96 recording solution comprising, in mM: KCl 2, NaCl 96, MgCl_2_ 1, and HEPES 5, buffered to pH 7.4 with KOH. Where indicated, K^+^ currents were assessed in a high-K^+^ recording solution comprising, in mM: KCl 96, NaCl 2, MgCl_2_1, and HEPES 5, buffered to pH 7.4 with KOH. Currents were measured from whole oocytes using an OC-725D amplifier (Warner Instruments), controlled via a USB interface (National Instruments) and were recorded using WinWCP software (University of Strathclyde). To evoke currents, oocytes were held at 0 mV, and currents were assessed by 160-ms ramps from -80 to +80 mV that were repeated every second. At the end of the study, currents were blocked by 6 mM BaCl_2_. The Ba^2+^-sensitive component of the current was analyzed. Between 4 and 12 oocytes from different *Xenopus* frogs were studied per experiment.

### Whole-cell patch-clamp recording

Whole-cell currents were recorded with a Tecella Pico-2 amplifier (Tecella) controlled using WinWCP software (Spider, University of Strathclyde). Currents were acquired through a low-pass Bessel filter at 2 kHz and were digitized at 10 kHz. Patch pipettes were fabricated from borosilicate glass (Clark Kent), using a vertical puller (Narishige) and had a resistance of 2.5–4 MΩ when filled with the intracellular buffers described below. HEK293T cells heterologously expressed the indicated channel subunits using a total of 1 µg DNA per transfection, with a DNA:PEI mass ratio of 1:4, and the subunits expressed at 1:1 DNA ratio unless otherwise indicated. Channel currents were studied using an intracellular buffer comprising, in mM: KCl 140, MgCl_2_ 2, EGTA 1, Na_2_ATP 5 and HEPES 5, adjusted to pH 7.2 with KOH. Following breakthrough into whole-cell mode, the cells were allowed to stabilize for 2 mins then currents were assessed by ramps from -80 to +80 mV that were repeated at 1 Hz. The external physiological recording buffer was applied via a gravity driven perfusion system. Cells were initially perfused with a low-K^+^ buffer comprising, in mM: NaCl 135, KCl 5, MgCl_2_ 1.2, CaCl_2_ 1.5, glucose 8, and HEPES 10, adjusted to pH 7.4 with NaOH, before being transitioned to a high-K^+^ buffer comprising, in mM: NaCl 5, KCl 135, MgCl_2_ 1.2, CaCl_2_ 1.5, glucose 8, and HEPES 10, adjusted to pH 7.4 with KOH. The Ba^2+^-sensitive component of the current was analyzed and determined by perfusing 3 mM BaCl_2_ in the high-K^+^ buffer at the end of each experiment. Cells for study were selected based on a fluorescence marker (typically eGFP) using an epifluorescence microscope (Olympus) equipped with patch-clamp. HEK293T cells had a mean whole-cell capacitance of 10 ± 8 pF, the series resistance was typically

<10 MΩ, and the voltage-error of 3 mV was not compensated.

### Calculation of residue decomposition binding energy

The binding energy between ML297 and its binding pocket was calculated based on the GIRK1/4-ML297 cryo-EM structure using the Discovery Studio program (Version 2017, Dassault Systèmes Biovia LLC, Waltham, MA). The channel complex was first energy-minimized with the Smart Minimizer algorithm (200 steps) using the CHARMm force field and the Generalized Born (GB) implicit solvent and membrane model. Interaction energies between the ligand and the channel were then computed using an implicit distance-dependent dielectric model (dielectric constant = 2.0). Residues within 6.5 Å of the ligand were included in the interaction-energy and residue-level energy-decomposition analyses.

### Gating Molecular Dynamic (GMD) Simulations

Molecular Dynamics simulations were performed on GIRK1/4^2:2^ -ML297(G1G4) and GIRK1/4^2:2^ -ML297(G1G1) complexes in Amber22 using ff19SB forcefield for proteins and Lipid21^65^ parameters for lipids. Potassium chloride (150mM) as well as neutralizing counterions were added to the systems. CHARMM-GUI Membrane Builder web server was used to embed complexes in a POPC : POPE : POPS : cholesterol (25:5:5:1) lipid bilayer. Systems were solvated in a rectangular TIP3P water box. The tleap module of Amber was used to generate coordinate and system topology files, followed by hydrogen-mass repartitioning (HMR) to enable a 3-fs integration timestep during production run. Particle-mesh Ewald (PME) and periodic boundary conditions were employed for long-range electrostatics. Energy minimization was performed by keeping the solute restrained with a harmonic force constant of 500 kcal mol⁻¹ Å⁻^2^ for 100 steps followed by 2000 steps of steepest descent. The system was heated gradually from 0 K to 100 K over 2500 steps using Langevin thermostat under constant volume while keeping proteins and lipids weakly restrained (500 kcal mol⁻¹ Å⁻^2^ for the solute; 10 kcal mol⁻¹ Å⁻^2^ for lipids). Subsequently, the temperature was gradually increased from 100 K to 303 K over 50,000 steps using anisotropic pressure coupling to reach 1 atm. System equilibration was done at 303 K under NPT conditions using Berendsen barostat and Langevin thermostat. SHAKE constraints were applied to all bonds and a time step of 1 fs was used. This was followed by a series of ten consecutive equilibration runs of 500 ps each (5 ns total) conducted at 303 K using a 2-fs time step. Weak restraints (10-20 kcal·mol⁻¹·Å⁻^2^) were applied to protein backbone to preserve stable membrane-protein packing during equilibration. Each of these equilibrated systems were subjected to 3 µs (3000 ns, six independent runs per system) of production run under NVT conditions, using 3fs time steps. A static electric field with the magnitude of 0.138 kcal·mol⁻¹·Å⁻^1^ (0.06 V/nm, 200mV across membrane) was applied along the negative z-direction. The results of the simulations were analyzed using various tools, including ChimeraX^57^, VMD^66^ and in-house scripts. GMD data files and details for others to replicate will become available through data bases depositing computational data.

### Interaction Pattern Analysis

CPPTRAJ and custom Python-based analysis scripts (written in pandas, numpy, and matplotlib) were utilized to explore key ligand-residue interactions throughout the simulation runs. Pairwise hydrogen-bond distances were calculated between critical protein residues and ligand donor/acceptor atoms. These included interactions involving Trp (W95.1), Glu (E141.1), Asp (D173.1), and Ser (S166.1, S176.4). The hydrogen-bond distances were monitored between ligand amide NH groups and carboxylate oxygens of Asp and Glu and ligand pyrazole nitrogen with serine hydroxyl protons. The contact was considered as hydrogen-bond when the donor-acceptor distance was ≤ 3.5 Å. Permeation events were also recorded and mapped with the interaction pattern data to compute total simulation time and frequency of ion permeations associated with each ligand interaction pattern.

## Data availability

The cryo-EM density maps and corresponding atomic coordinates have been deposited in the Electron Microscopy Data Bank (EMDB) and the PDB, respectively, under the following accession codes: EMD-74952 and 9ZYG (GIRK1/4^1:3^); EMD-74953 and 9ZYH (GIRK1/4^2:2^); EMD-74954 and 9ZYI (ML297-bound GIRK1/4^1:3^ complex); and EMD-74955 and 9ZYJ (ML297-bound GIRK1/4^2:2^(G1G1) complex); EMD-74956 and 9ZYK (ML297-bound GIRK1/4^2:2^(G1G4) complex); EMD-74957 and 9ZYL (GIRK4). Source data are provided with this paper.

## Acknowledgements

This work was supported by NIH R01 grants HL059949-28 and NS131467-03 to DEL. Thanks to: Heikki Vaananen for EP technical guidance and admin support, the following colleagues for feedback on the manuscript: Stelios Smirnakis and Mahmoud Nasr at Brigham & Women’s Hospital at Harvard Medical School, Sami Noujaim at the University of South Florida and Satya Kunapuli at Temple University, Logothetis/Plant lab members for discussions. The computations were supported by the ITS (Information Technology Services) Research Computing at Northeastern University and the Argonne Leadership Computing Facility (ALCF) at Argonne National Laboratory.

## Author contributions

L.W., T.K., D.E.L., G.S., L.P., and M.C. designed the study. L.W. performed cryo-EM sample preparation, data collection, processing, model building, structural analysis. T.K. purified proteins, prepared all mutants, with assistance from N.A. L.P. and J.P. performed single-molecule TIRF experiments. A.S.-C. and E.D. carried out patch-clamp and A.Z. and K.W.C. TEVC recordings. N.P., M.Z. and M.C. supervised all computational studies. D.E.L. and G.S. conceived and supervised the study. L.W. and L.D.P. prepared the figures. L.W., D.E.L. and G.S. wrote the manuscript with input from all authors.

