## SupplementaryData for "Stoichiometry-dependent gating and structural pharmacology of cardiac GIRK1/4 channels"

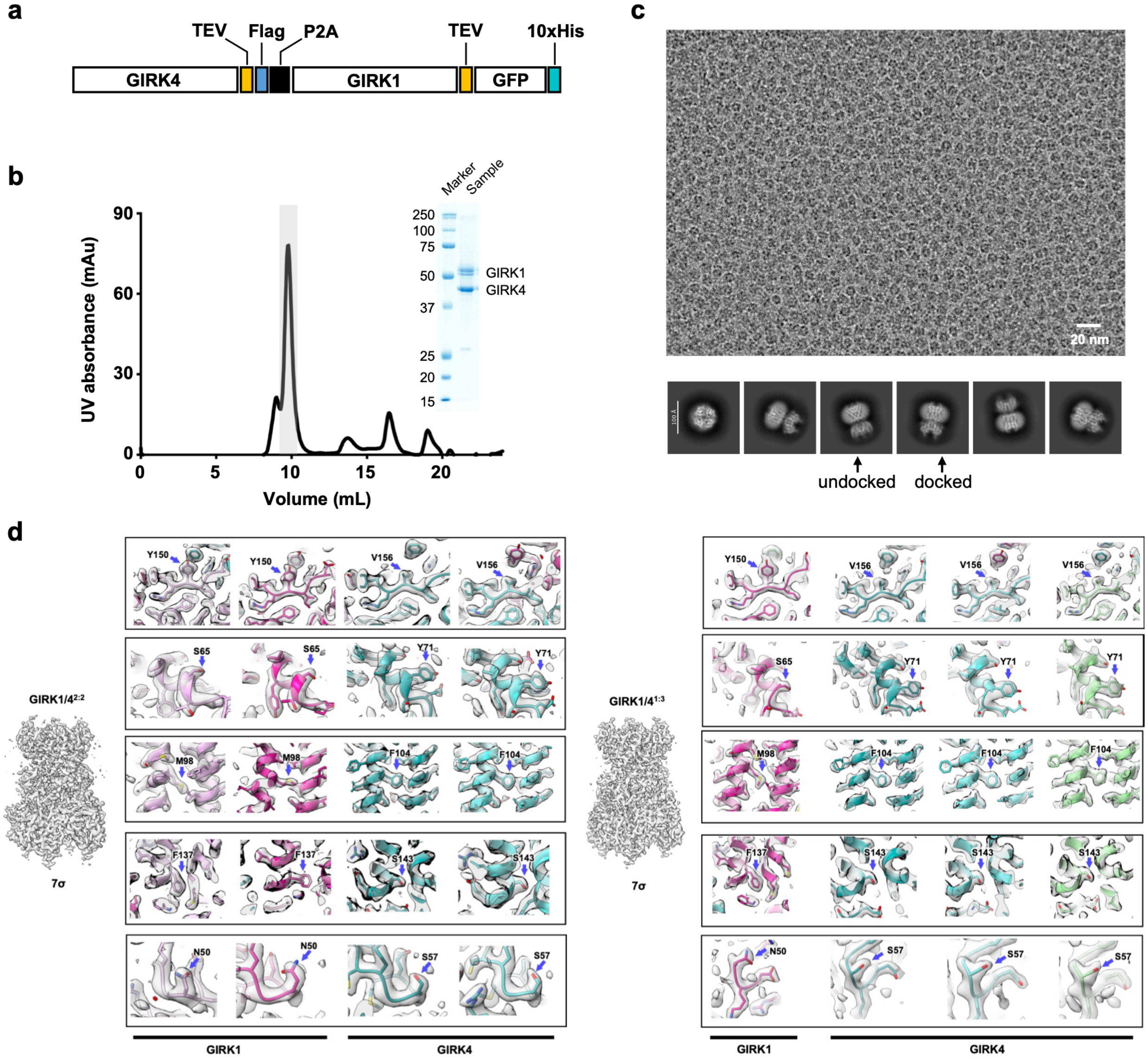


**Extended Data Fig. 1 | Expression, purification, and cryo-EM analysis of the GIRK1/4 heterotetramer. a**, Schematic of the GIRK4-P2A-GIRK1-GFP construct used for co-expression of GIRK4 and GIRK1 subunits. TEV-cleavable sites were inserted between each gene and its adjacent tag. **b**, Size-exclusion chromatography of the purified GIRK1/4 complex showing a major tetramer peak (shaded). Inset, SDS-PAGE confirming the presence of GIRK1 and GIRK4. **c**, Representative cryo-EM micrograph and selected 2D class averages showing undocked and docked conformations of the CTD relative to the TMD. **d**, Representative cryo-EM densities for residue-specific features used to assign GIRK1 and GIRK4 subunits in the GIRK1/4$^{1:3}$ and GIRK1/4$^{2:2}$ maps. GIRK1 is shown in magenta, and GIRK4 subunits are shown in teal/green. The highlighted residues show subunit-specific density features that distinguish GIRK1 from GIRK4 and support assignment of the asymmetric heterotetrameric maps. Grey mesh represents the cryo-EM density. All maps are displayed at a contour level of 7σ.

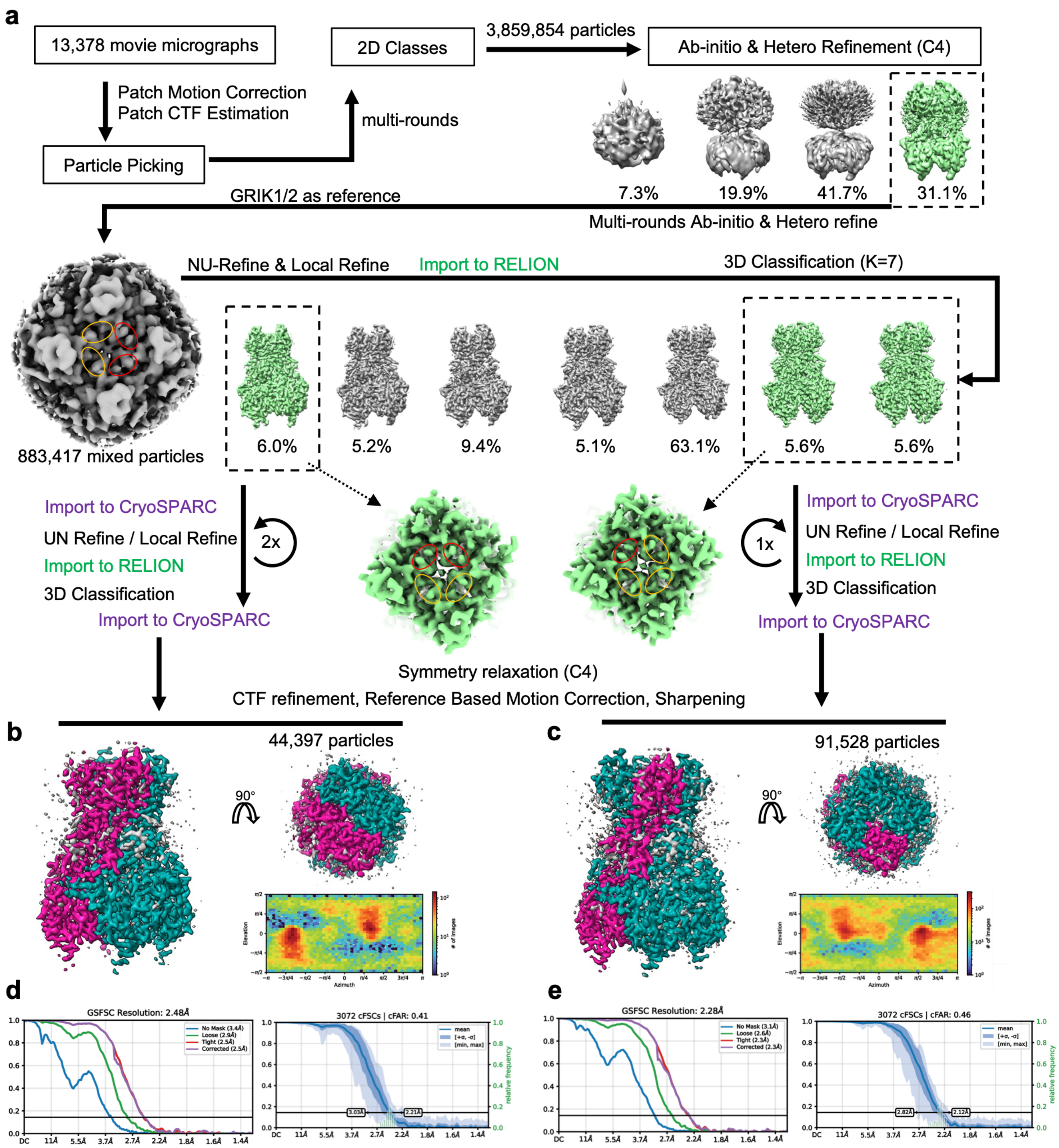


**Extended Data Fig. 2 | Cryo-EM processing and reconstruction of GIRK1/4 in two different stoichiometries.** **a**, Cryo-EM data processing workflow for the GIRK1/4 complex in docked state. **b**, Reconstruction of GIRK1/4$^{2:2}$ (GIRK1 in red and GIRK4 in teal) and its angular distribution heat map of particle projections. **c**, Reconstruction of GIRK1/4$^{1:3}$ and its angular distribution heat map of particle projections. **d**, The Gold-standard Fourier shell correlation (GSFSC) curve (left) indicates a global resolution of 2.48 Å at the 0.143 cutoff. The conical FSC (cFSC) plot (right) shows the mean (dark line) and variation (blue shading) of directional correlations, with a calculated cFAR (conical FSC Area Ratio) of 0.41, indicating modest directional anisotropy of the map. **e**, The GSFSC curve (left) indicates a global resolution of 2.28 Å at the 0.143 cutoff. The cFSC plot (right) shows the mean and variation of directional correlations, with a calculated cFAR of 0.46.

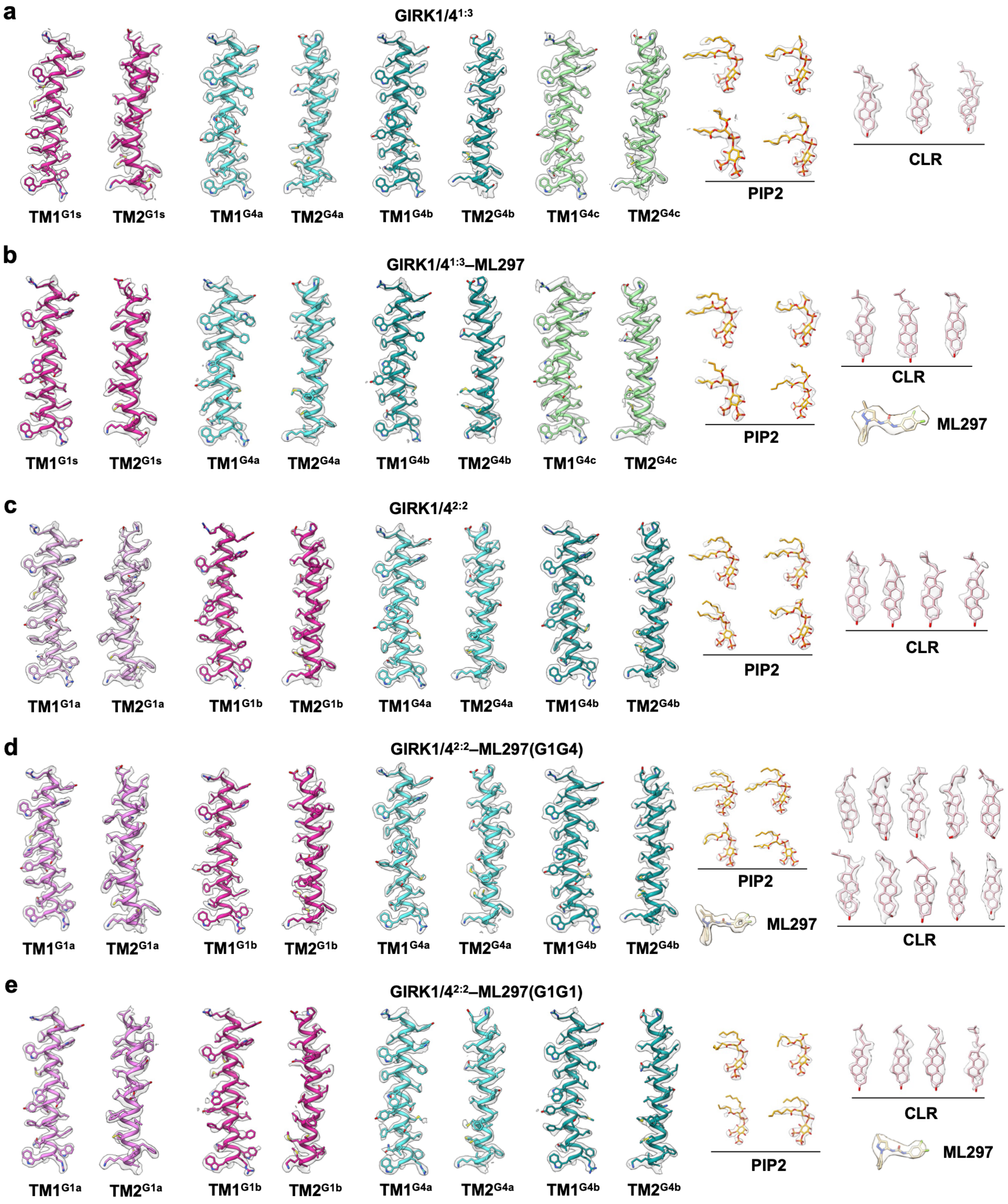


**Extended Data Fig. 3 | Agreement between cryo-EM density and model. a**, EM density and model for TM1, TM2, $PIP_2$ and Cholesterol in the GIRK1/$4^{1:3}$ structure. **b**, EM density and model for TM1, TM2, $PIP_2$ and Cholesterol in the ML297-bound GIRK1/$4^{1:3}$ structure. **c**, EM density and model for TM1, TM2, $PIP_2$ and Cholesterol in the GIRK1/$4^{2:2}$ structure. **d**, EM density and model for TM1, TM2, $PIP_2$ and Cholesterol in the ML297-bound GIRK1/$4^{2:2}$ structure, in which ML297 binds to G1G4 pocket. **e**, EM density and model for TM1, TM2, $PIP_2$ and Cholesterol in the ML297-bound GIRK1/$4^{2:2}$ structure, in which ML297 binds to G1G1 pocket.

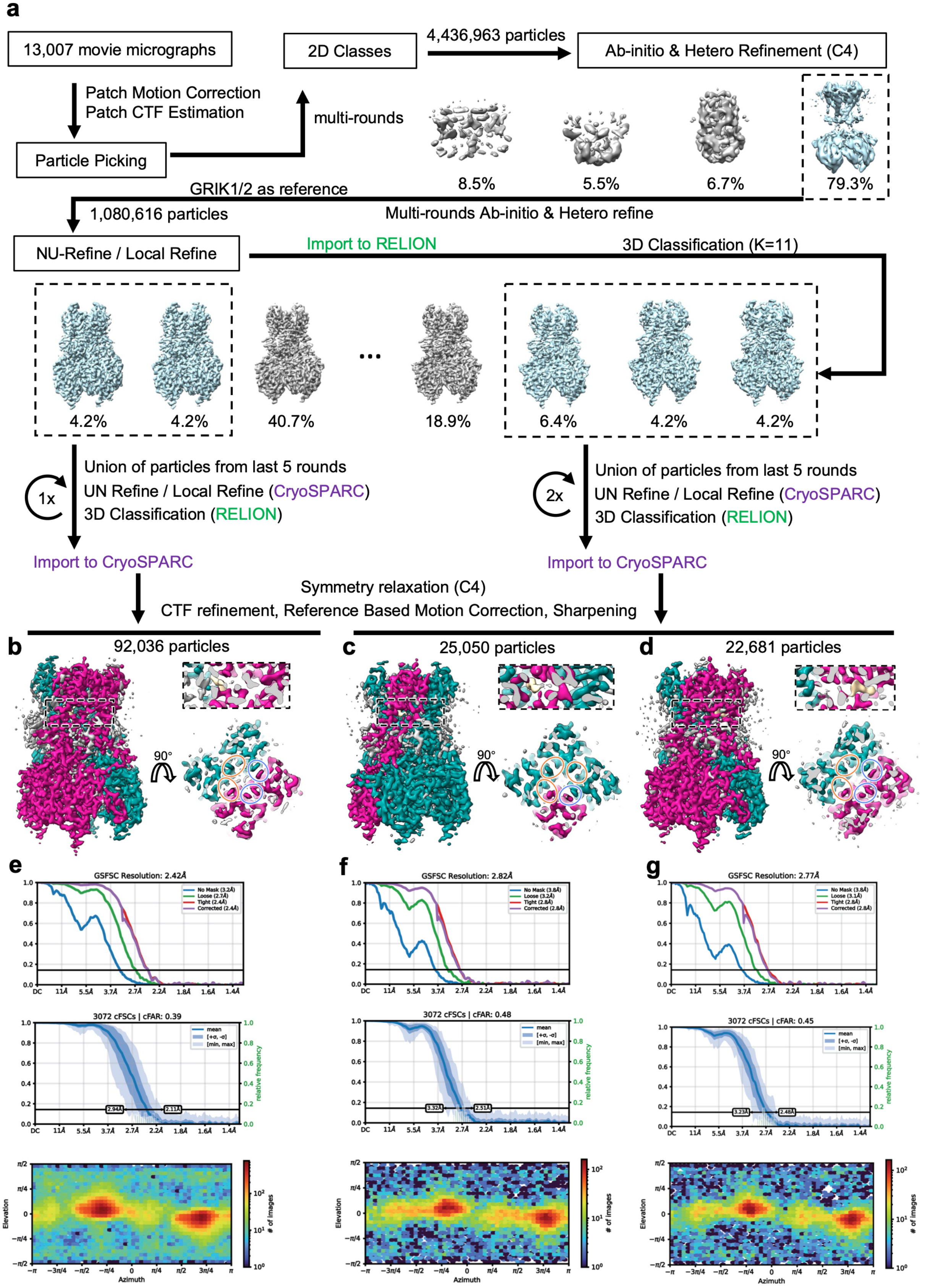
a
13,007 movie micrographs
Patch Motion Correction
Patch CTF Estimation
Particle Picking
multi-rounds
2D Classes
4,436,963 particles
Ab-initio & Hetero Refinement (C4)
8.5%
5.5%
6.7%
79.3%
GRIK1/2 as reference
Multi-rounds Ab-initio & Hetero refine
1,080,616 particles
NU-Refine / Local Refine
Import to RELION
3D Classification (K=11)
4.2%
4.2%
40.7%
...
18.9%
6.4%
4.2%
4.2%
1x
Union of particles from last 5 rounds
UN Refine / Local Refine (CryoSPARC)
3D Classification (RELION)
Import to CryoSPARC
2x
Union of particles from last 5 rounds
UN Refine / Local Refine (CryoSPARC)
3D Classification (RELION)
Import to CryoSPARC
Symmetry relaxation (C4)
CTF refinement, Reference Based Motion Correction, Sharpening
b
92,036 particles
90°
c
25,050 particles
90°
d
22,681 particles
90°
e
GSFSC Resolution: 2.42Å
f
GSFSC Resolution: 2.82Å
g
GSFSC Resolution: 2.77Å
3072 cFSCs | cFAR: 0.39
3072 cFSCs | cFAR: 0.48
3072 cFSCs | cFAR: 0.45
Elevation
Azimuth
### of images

**Extended Data Fig. 4 | Cryo-EM processing and reconstruction of ML297-bound GIRK1/4 in different stoichiometries.** **a**, Cryo-EM data processing workflow for the ML297-bound GIRK1/4 complex. **b**-**d**, Reconstructions of GIRK1/$4^{2:2}$-ML297(G1G4 pocket) (**b**), GIRK1/$4^{1:3}$-ML297 (**c**), and GIRK1/$4^{2:2}$-ML297(G1G1 pocket) (**d**). GIRK1 and GIRK4 are colored red and teal, respectively. Boxed regions highlight the ML297-binding pockets, and circles mark structural features that differ between GIRK1 and GIRK4. **e**-**g**, GSFSC curves, cFSC plots, and angular distribution heat maps of particle projections corresponding to the reconstructions of GIRK1/$4^{2:2}$-ML297(G1G4 pocket) (**e**), GIRK1/$4^{1:3}$-ML297 (**f**), and GIRK1/$4^{2:2}$-ML297(G1G1 pocket) (**g**).

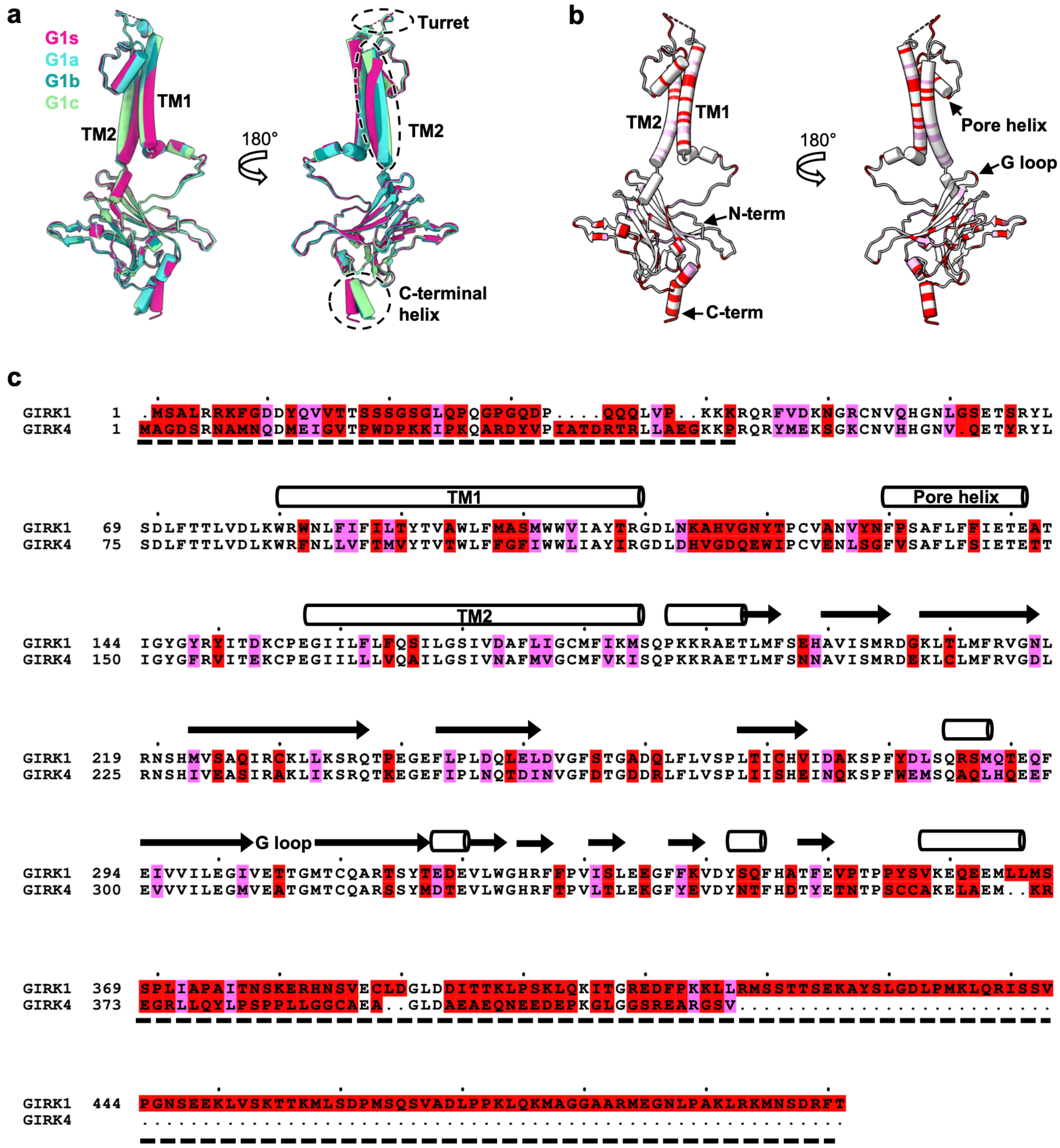

**Extended Data Fig. 5 | Structure and sequence comparison between GIRK1 and GIRK4 subunits.** **a**, Structural comparison of individual subunits from GIRK1/4[1:3] heterotetramer showing conformational variations among GIRK1 and GIRK4 subunits. Dashed ovals highlight regions with pronounced differences. **b**, Structural mapping of sequence divergence between GIRK1 and GIRK4. Red, purple, and white indicate residues that are different, similar, or identical, respectively. **c**, Sequence alignment of GIRK1 and GIRK4. Residues with large differences are colored red, and similar residues are purple, corresponding to the coloring in panel (**b**). Secondary structure elements (TM1, TM2, pore helix, and G loop) are indicated above the alignment. Thick dashed lines mark unresolved regions at the N- and C-termini in the cryo-EM density map.

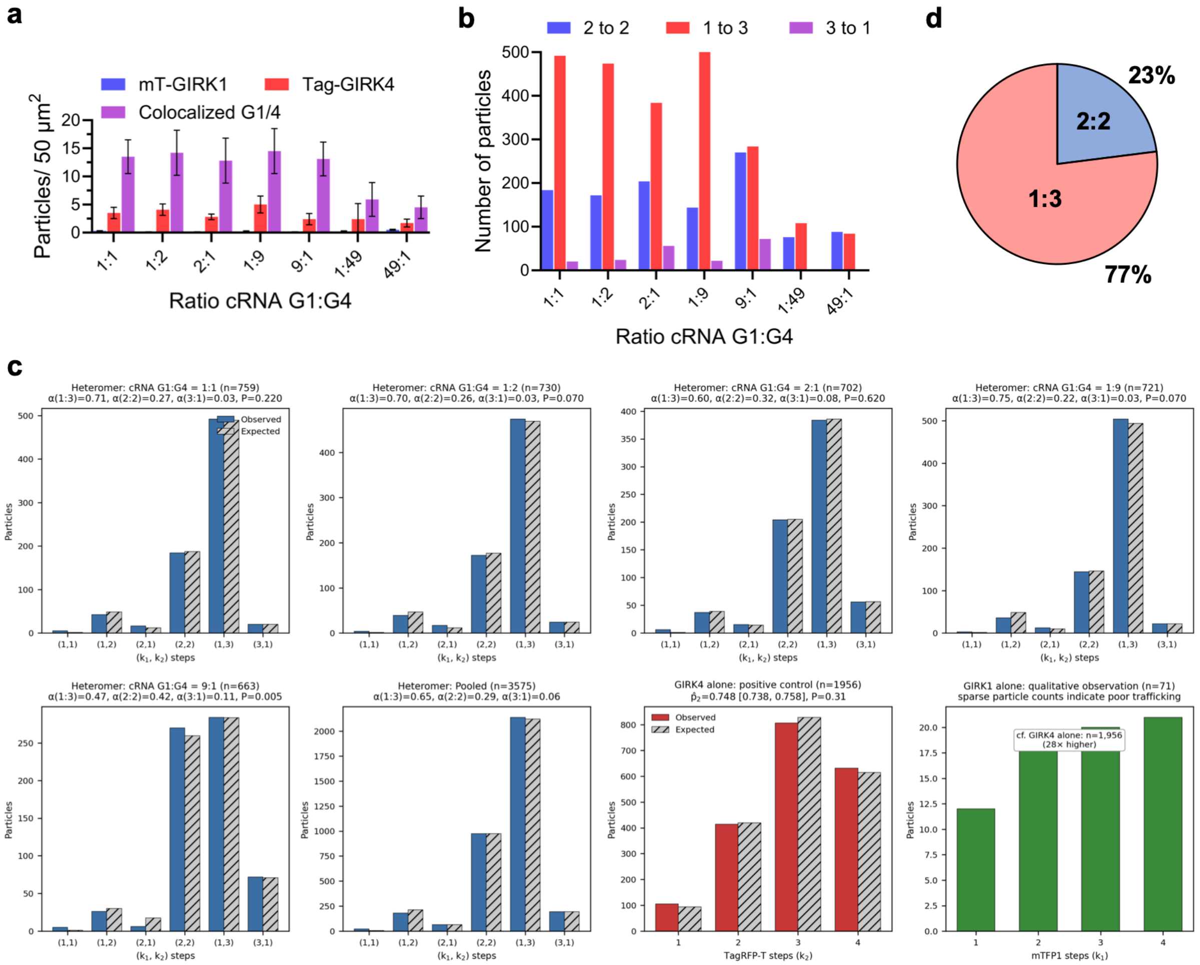


**Extended Data Fig. 6 | Stoichiometry distribution of GIRK1/4 heterotetramers under varying expression ratios.** *Xenopus* oocytes were co-injected to express mTFP1-tagged (mT)-GIRK1 and TagRPT-T (Tag)-GIRK4 at the ratios indicated. Fluorescent particles were visualized by TIRF microscopy as mT-GIRK1 alone (blue), Tag-GIRK4 alone (red) or colocalized (purple). Data are from 8-10 oocytes per group from at least 3 independent replicates of the experiment. **a**, Summary histogram of the number of individual or colocalized particles counted per 50 $\mu m^2$ at the GIRK1 : GIRK4 injection ratio indicted. **b**, Summary histogram showing the distribution of subunit compositions for colocalized particles counted for each injection ratio, displayed in panel A. Particles had three apparent stoichiometries: GIRK1 to GIRK4 at 2:2 (blue), 1:3 (red), or 3:1 (purple). **c**, Binomial photobleaching fits for colocalized GIRK1/4 particles across five GIRK1 cRNA injection ratios and for the pooled heteromer dataset. Observed bleaching-step counts are shown as solid bars and model-expected counts are shown as hatched bars. Each heteromer particle was assigned by the number of resolved mTFP1 and TagRFP-T bleaching steps, denoted as $(k_1, k_2)$, where $k_1$ corresponds to GIRK1-mTFP1 and $k_2$ corresponds to GIRK4-TagRFP-T. The fitted model included three GIRK1 stoichiometries, 1:3, 2:2 and 3:1, with per-fluorophore detection probabilities shared across conditions. GIRK4 expressed alone was fit as a positive-control tetrameric population, whereas GIRK1 expressed alone is shown as a qualitative trafficking control because of sparse membrane particle counts. **d**, Distribution of particle populations from cryo-EM classification of GIRK1/4 samples, with 77% corresponding to GIRK1/$4^{1:3}$ and 23% to GIRK1/$4^{2:2}$ stoichiometries. Statistics from cryo-EM data after removing the mixed particles.

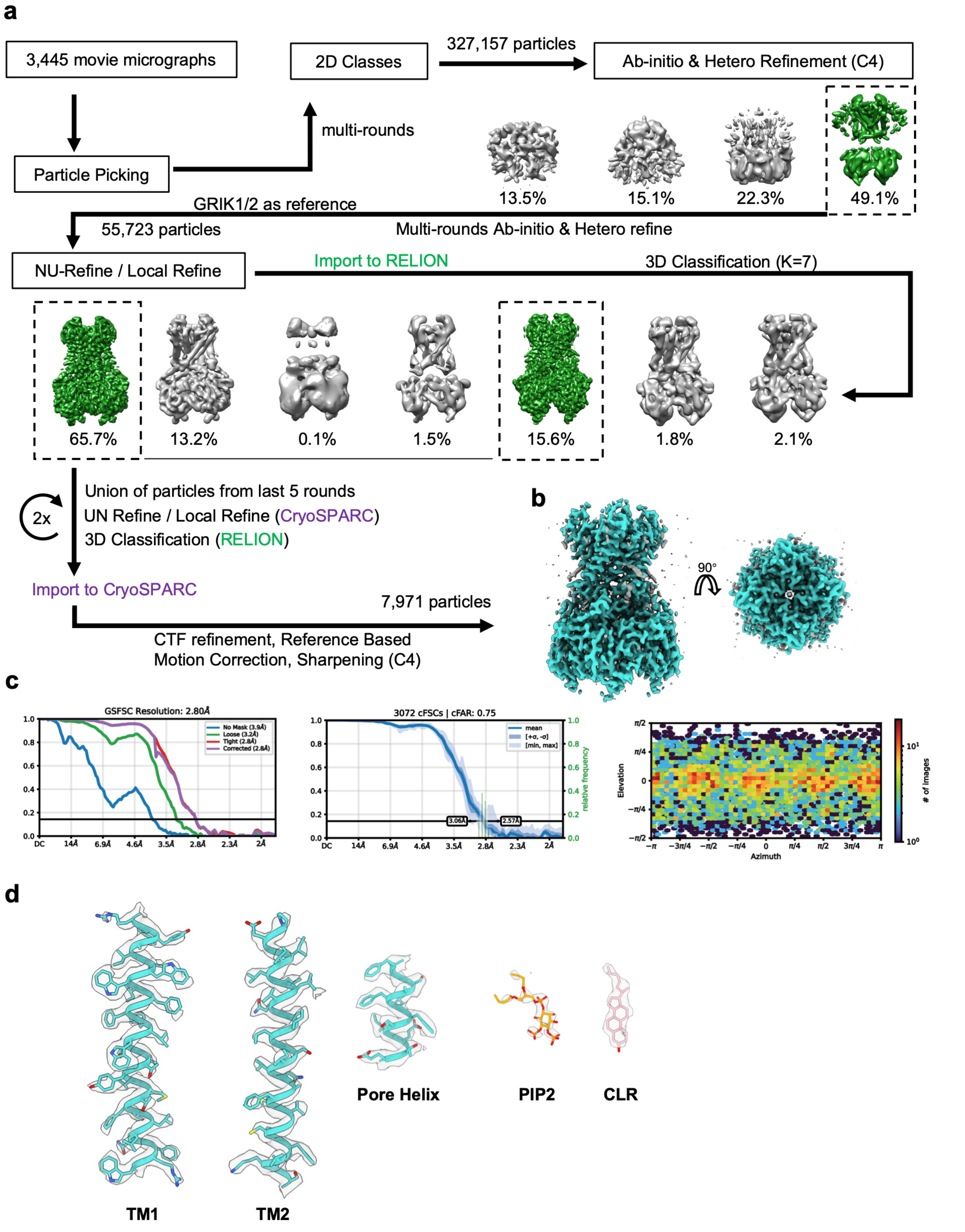


**Extended Data Fig. 7 | Cryo-EM processing and reconstruction of GIRK4 homotetramer. a**, Cryo-EM data processing workflow for the GIRK4 complex. **b**, Reconstructions of GIRK4. **c**, GSFSC curves, cFSC plots, and angular distribution heat maps of particle projections. **d**, EM density and model for TM1, TM2, pore helix, $PIP_2$ and Cholesterol in the GIRK4 structure.

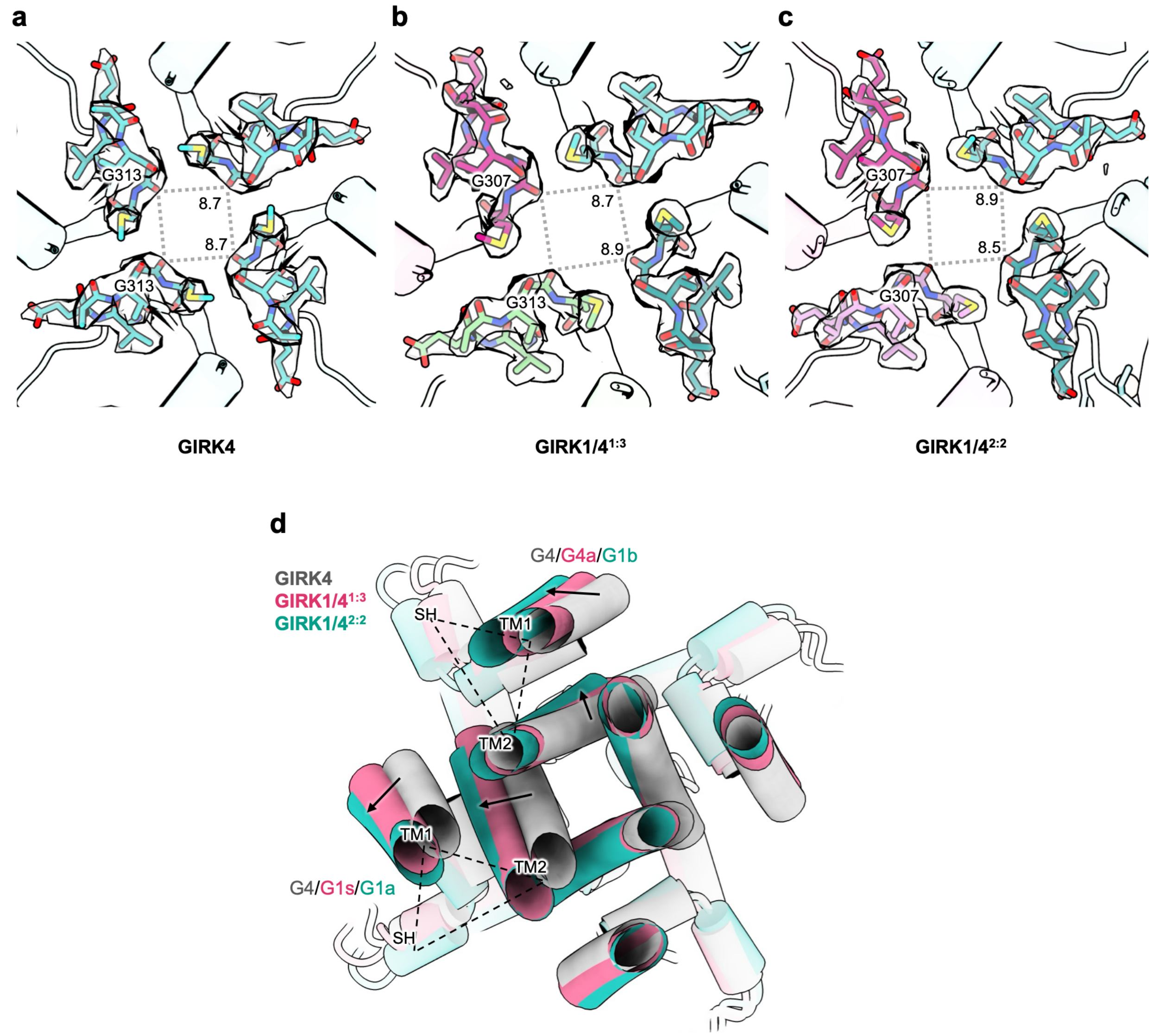


**Extended Data Fig. 8 | Structure comparison of the G-loop gate and HBC opening among GIRK4 and GIRK1/4 heterotetramers. a-c**, Cryo-EM densities and atomic models of the G-loop gate in GIRK4 (**a**), GIRK1/4$^{1:3}$ (**b**), and GIRK1/4$^{2:2}$ (**c**). The backbone carbonyls of glycine residues (Gly313 in GIRK4, Gly307 in GIRK1) define the narrowest constriction of the G-loop gate. The values indicate the shortest distances between opposing subunits that define these constrictions. **d**, Supplementary view to Fig. 2f, showing a slice of the superimposed GIRK4, GIRK1/4$^{1:3}$, and GIRK1/4$^{2:2}$ structures. This view more clearly illustrates the progressive widening of the TMD bottom near the HBC as GIRK1 subunits are incorporated.

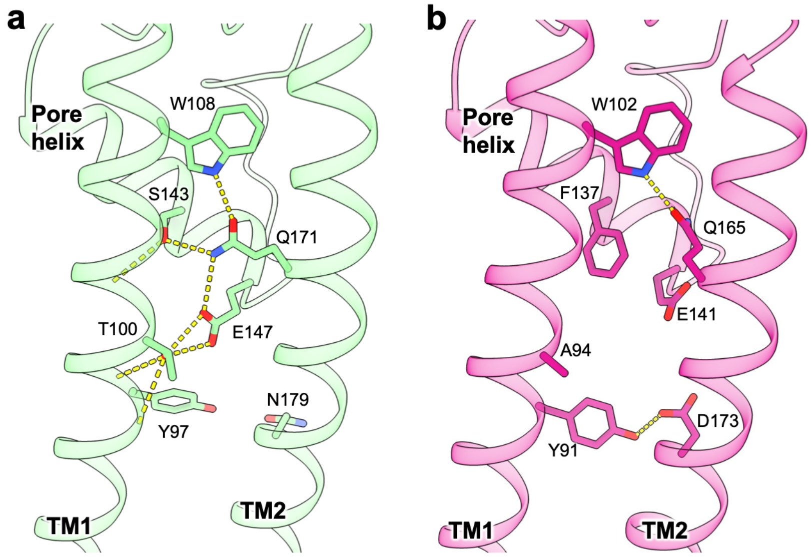


**Extended Data Fig. 9 | Comparison of intra-subunit hydrogen-bond networks between GIRK4 and GIRK1. a**, Hydrogen-bond network in the GIRK4 subunit connecting residues from TM1, TM2 and the pore helix. **b**, Corresponding view of the GIRK1 subunit, showing a distinct pattern of polar interactions between TM1 and TM2. Hydrogen bonds are shown as yellow dashed lines.

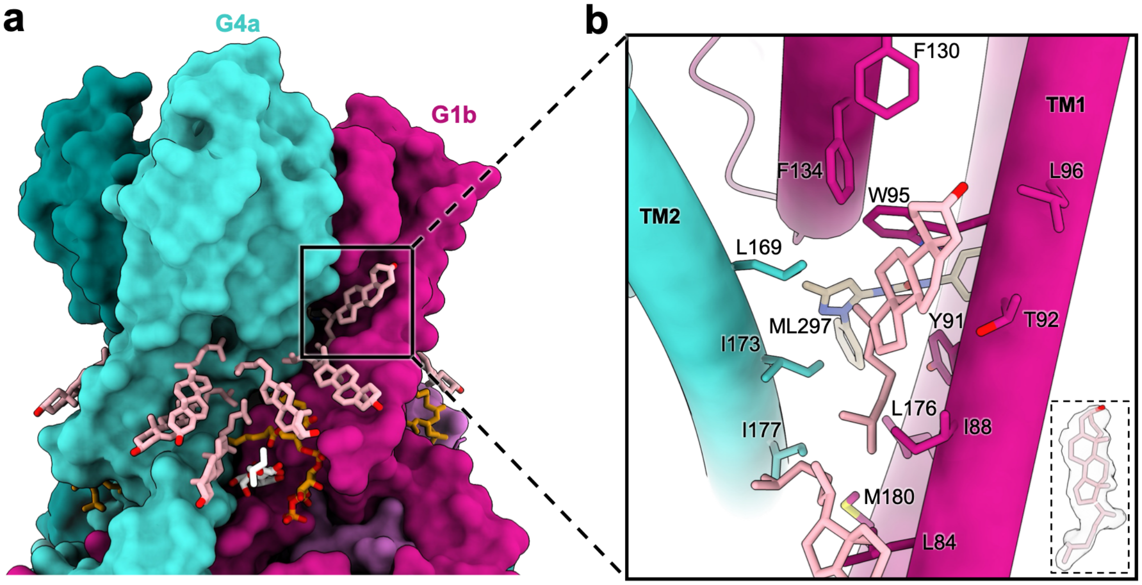


**Extended Data Fig. 10 | Cholesterol binding at the GIRK1-GIRK4 interface. a**, Surface representation of the GIRK1/$4^{2:2}$-ML297(G1G4) heterotetramer showing the cholesterol-binding pocket located at the interface between GIRK1 and GIRK4 subunits. **b**, Close-up view of the boxed region in (**a**), highlighting hydrophobic interactions that stabilize cholesterol binding. Key residues from both subunits are labeled, and the inset shows the fitted density for cholesterol.

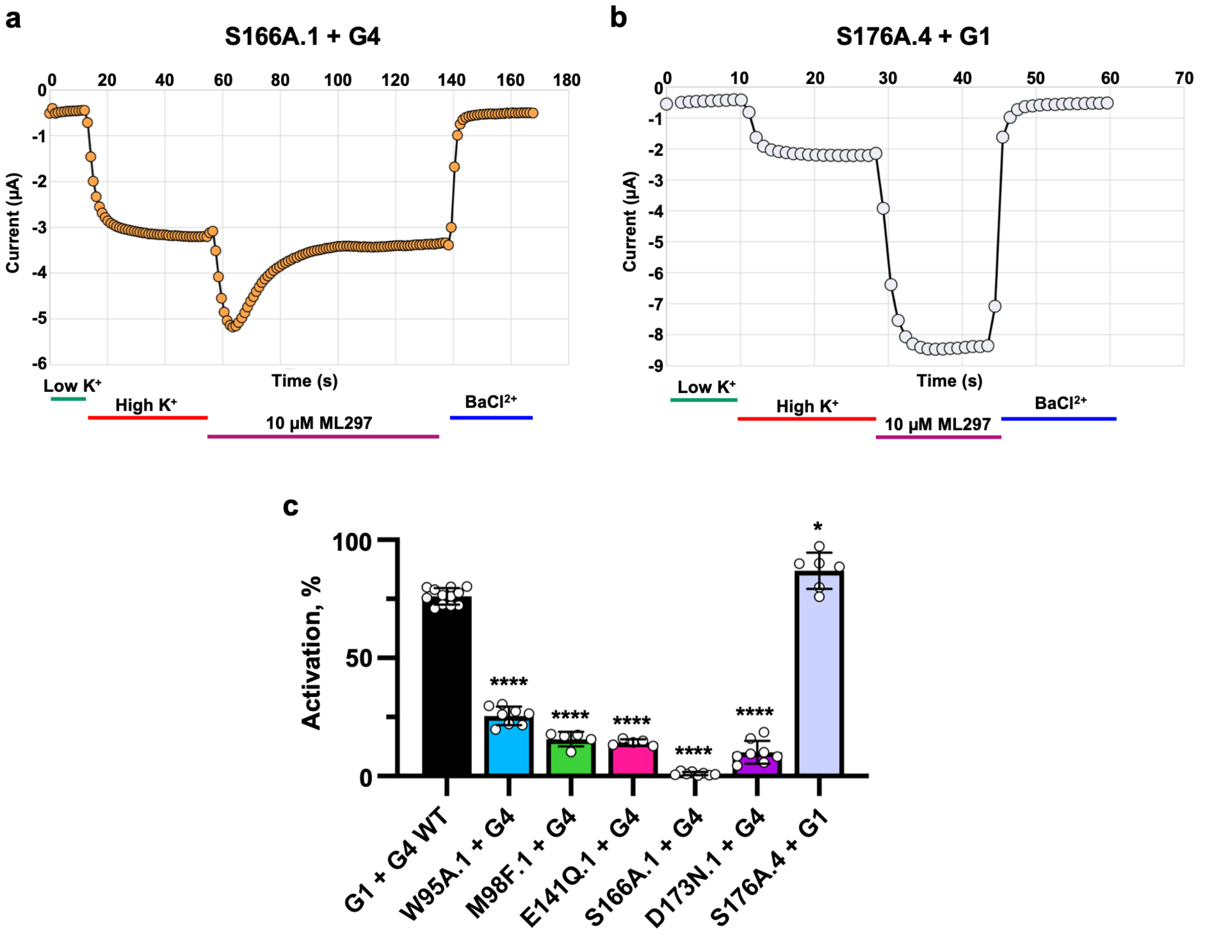


**Extended Data Fig. 11 | Examples of mutants with most extreme effects on ML297 activation in oocytes and comparison of mutant effects in mammalian cells. a**,**b**, Representative traces from TEVC recordings in *Xenopus* oocytes showcasing the effect of ML297 on the S166A.1 (**a**) and S176C.4 (**b**) mutants, where steady-state currents are abolished in S166A.1 but enhanced in S176C.4. **c**, Summary %-current activation in response to 10 mM ML297 from HEK-293 cells expressing GIRK1 + GIRK4 with the mutations indicated, in the GIRK1 or the GIRK4 subunit. Statistical significance is assessed compared to the wild type (WT) subunits. The inset shows example current sweeps from the WT channel. Whole-cell patch-clamp was performed as described in the Methods. Summary data are from 6-10 cells per group from 3 independent experiments. Statistical significance was assessed using a two-tailed, unpaired Welches T-test where *P<0.05 and ****P<0.0001.

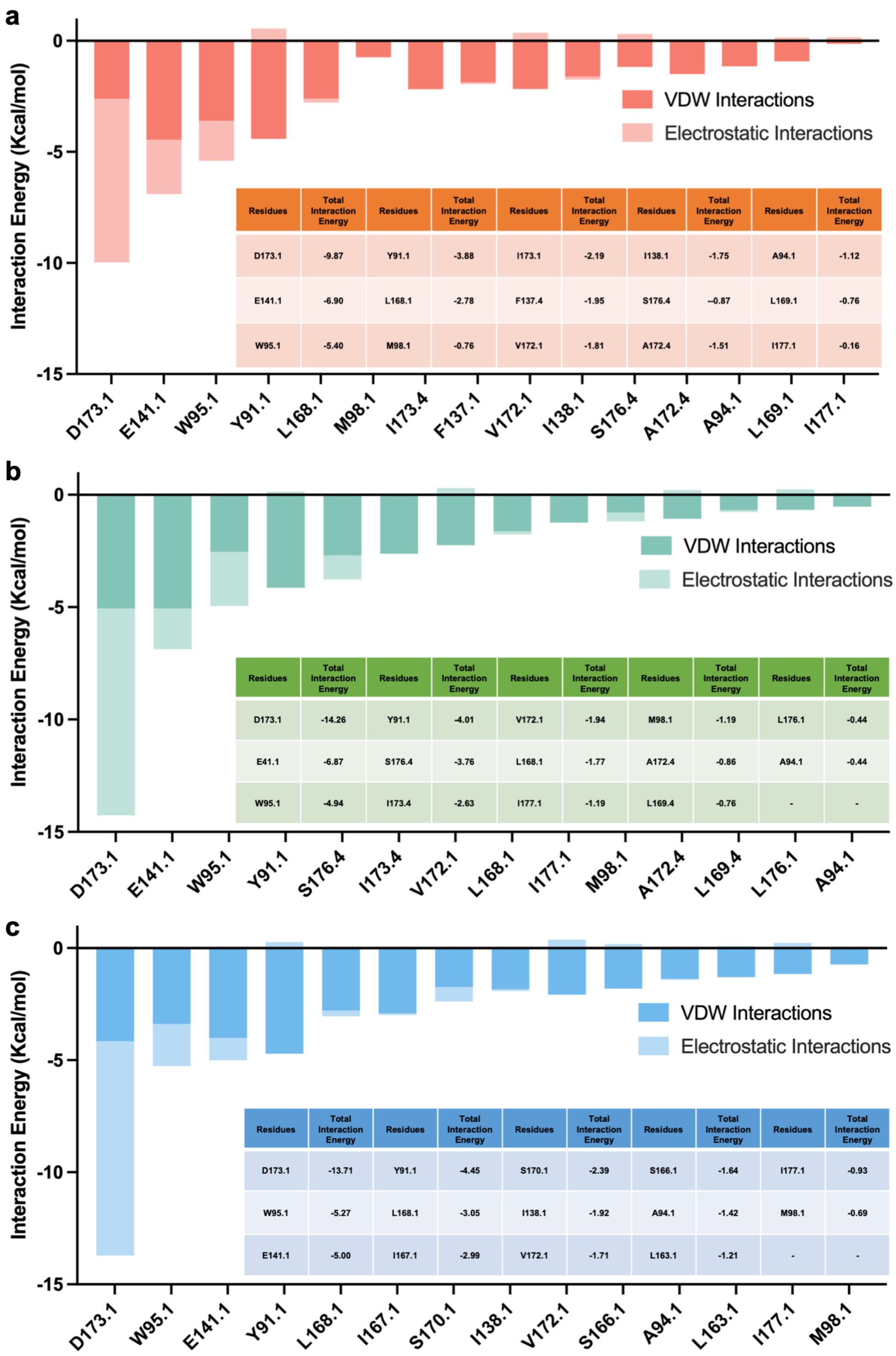


**a**

| Residues | Total Interaction Energy | Residues | Total Interaction Energy | Residues | Total Interaction Energy | Residues | Total Interaction Energy | Residues | Total Interaction Energy |
|---|---|---|---|---|---|---|---|---|---|
| D173.1 | -9.87 | Y91.1 | -3.88 | I173.1 | -2.19 | I138.1 | -1.75 | A94.1 | -1.12 |
| E141.1 | -6.90 | L168.1 | -2.78 | F137.4 | -1.95 | S176.4 | --0.87 | L169.1 | -0.76 |
| W95.1 | -5.40 | M98.1 | -0.76 | V172.1 | -1.81 | A172.4 | -1.51 | I177.1 | -0.16 |

**b**

| Residues | Total Interaction Energy | Residues | Total Interaction Energy | Residues | Total Interaction Energy | Residues | Total Interaction Energy | Residues | Total Interaction Energy |
|---|---|---|---|---|---|---|---|---|---|
| D173.1 | -14.26 | Y91.1 | -4.01 | V172.1 | -1.94 | M98.1 | -1.19 | L176.1 | -0.44 |
| E41.1 | -6.87 | S176.4 | -3.76 | L168.1 | -1.77 | A172.4 | -0.86 | A94.1 | -0.44 |
| W95.1 | -4.94 | I173.4 | -2.63 | I177.1 | -1.19 | L169.4 | -0.76 | - | - |

**c**

| Residues | Total Interaction Energy | Residues | Total Interaction Energy | Residues | Total Interaction Energy | Residues | Total Interaction Energy | Residues | Total Interaction Energy |
|---|---|---|---|---|---|---|---|---|---|
| D173.1 | -13.71 | Y91.1 | -4.45 | S170.1 | -2.39 | S166.1 | -1.64 | I177.1 | -0.93 |
| W95.1 | -5.27 | L168.1 | -3.05 | I138.1 | -1.92 | A94.1 | -1.42 | M98.1 | -0.69 |
| E141.1 | -5.00 | I167.1 | -2.99 | V172.1 | -1.71 | L163.1 | -1.21 | - | - |

**Extended Data Fig. 12 | Per-residue decomposition of ML297 interaction energies in different GIRK1/4 stoichiometries and binding pockets.** Decomposed interaction energy profiles showing the contribution of individual residues to ML297 binding in various GIRK1/4 stoichiometries and binding poses**.** Bar graphs display van der Waals (VDW) and electrostatic interaction energy components for each residue. Embedded tables provide the total interaction energy (sum of VDW and electrostatic contributions) for each residue. **a**, ML297 interaction energy decomposition in the GIRK $14^{1:3}$. **b**, ML297 interaction energy decomposition in the GIRK $14^{2:2}$(G1G4) pocket. **c**, ML297 interaction energy decomposition in the GIRK $14^{2:2}$(G1G1) pocket.

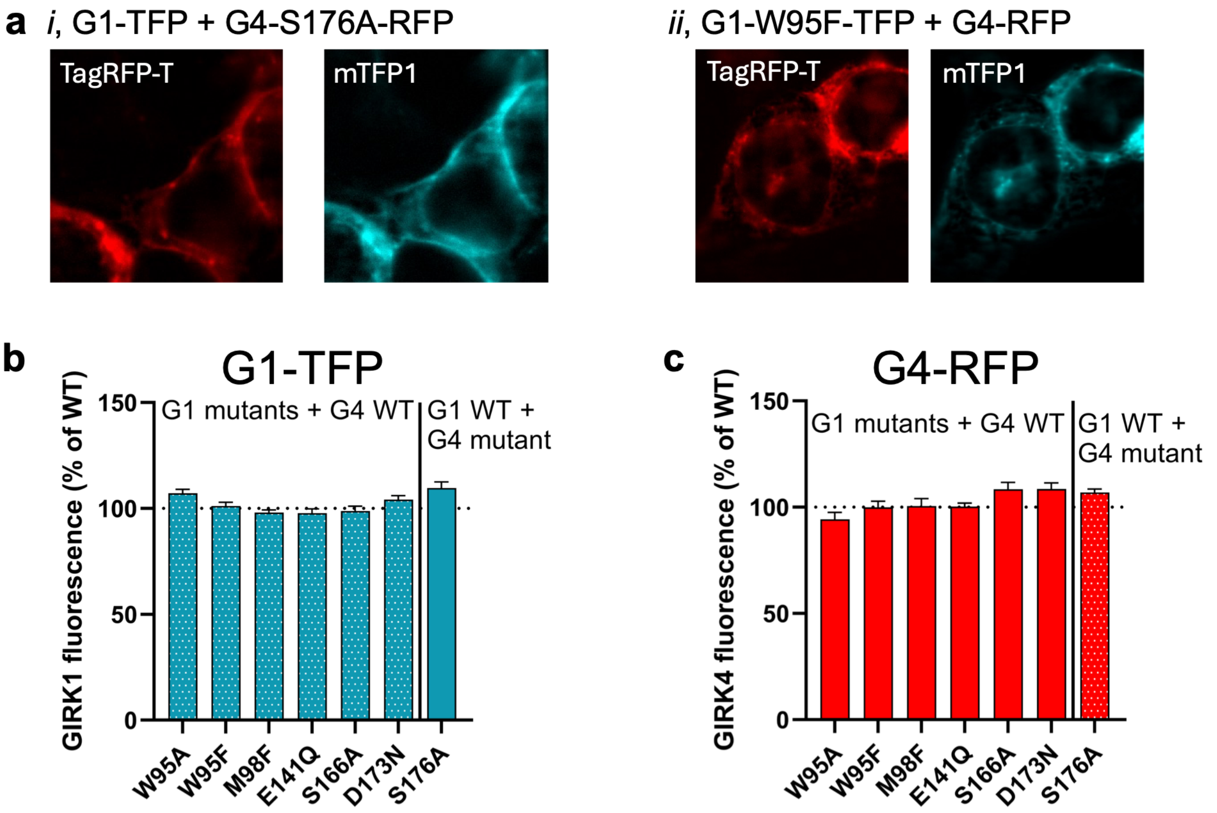


**Extended Data Fig. 13 | Cell surface localization of GIRK1 or GIRK4 mutants co-expressed with the complementary heterotetrameric subunit.**

HEK293T cells imaged by TIRF microscopy following transfection of the indicated GIRK1-TFP and GIRK4-RFP variants. The mean fluorescence intensities of both fluorophores were measured for all conditions and are presented as mean ± s.e.m. relative to wild-type control from 15-25 cells per condition across 3 biological replicates. For both fluorophores, fluorescence relative to GIRK1 wild-type + GIRK4 wild-type condition was consistent across all constructs. **a**, Representative fluorescence micrographs. **b**, Summary histogram of GIRK1-TFP fluorescence in the GIRK1/4 variants indicated. **c**, Summary histogram of GIRK4-RFP fluorescence in the GIRK1/4 variants indicated.

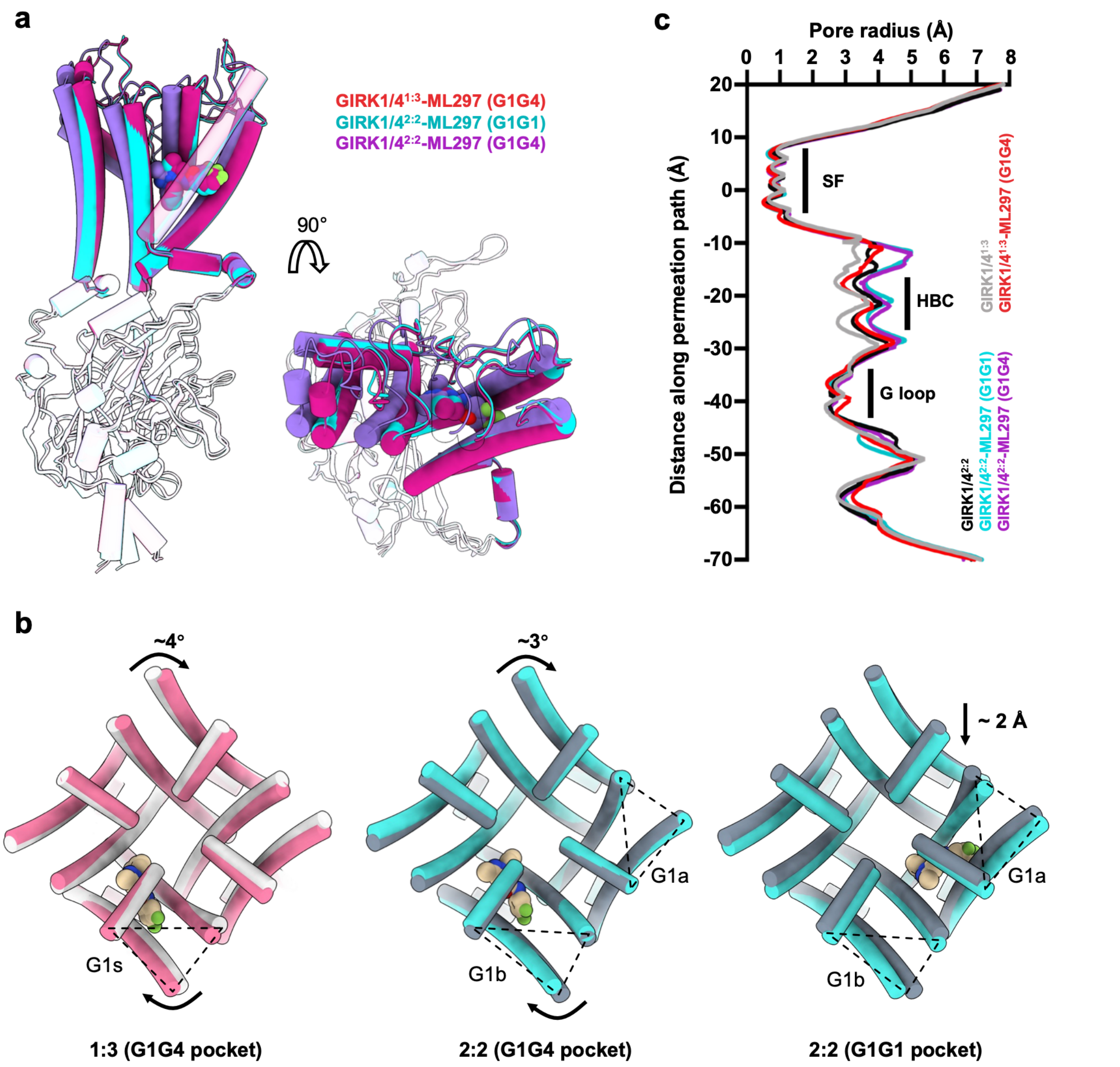


**Extended Data Fig. 14 | Structural comparison of ML297 binding and pore architecture in GIRK1/4 heterotetramers. a**, Superposition of the ML297-bound GIRK1/4 structures highlighting different binding interfaces. Side and top views show distinct subunit arrangements associated with each binding mode. **b**, Comparison between apo (without ML297) and ML297-bound GIRK1/4 structures viewed from the extracellular side. **c**, Pore-radius profiles calculated using HOLE, comparing the permeation pathways among five GIRK1/4 structures in this study. For the ML297-bound GIRK1/4$^{1:3}$ structure, the red trace represents the pore-radius profile measured from the conformation with the largest HBC size.

**a**

| SYSTEMS | Interactions involving residue E141.1/D173.1/W95.1/S166.1/S176.4 | | Total ST (μs) | Total PC |
|---|---|---|---|---|
| | ST in *ns* | PC | | |
| ML-G1/4$^{1:3}$ (G1G4) | E: 0<br>D: 1906 (63.5%)<br>W: 797.8 (26.9%)<br>S176: 389 (13%) | E: 0<br>D: 44 (26%)<br>W: 39 (23.1%)<br>S176: 17 (10.1 %) | 3 | 169 |
| WT-G1/4$^{1:3}$ | - | - | | 208 |
| ML-G1/4$^{2:2}$ (G1G1) | E: 0<br>D: 2299 (77%)<br>W: 2073 (70.3%)<br>S166 : 1377(46%) | E : 0<br>D: 178 (67%)<br>W: 206 (78%)<br>S166: 132 (50%) | | 264 |
| ML-G1/4$^{2:2}$ (G1G4)<br>-CHL | E: 482 (16.2%)<br>D: 2622 (87.5%)<br>W: 1831 (60.9%)<br>S: 978(32.7%) | E: 22 (17.3%)<br>D: 115 (89.5%)<br>W: 75 (58.4%)<br>S: 40 (31.2%) | | 129 |
| ML-G1/4$^{2:2}$ (G1G4)<br>+CHL | E: 93 (3.4%)<br>D: 2996 (99.8%)<br>W: 2992 (99.7%)<br>S: 697.6 (23.5 %) | E: 2 (1%)<br>D: 201 (100%)<br>W: 201 (100%)<br>S: 91 (45.3%) | | 201 |
| WT-G1/4$^{2:2}$ | - | - | | 157 |

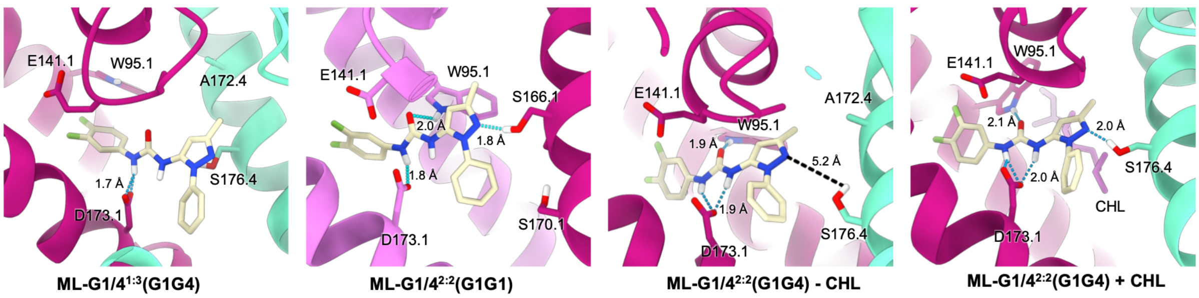


**Extended Data Fig. 15 | GMD simulations on GIRK1/4$^{1:3}$-ML297, GIRK1/4$^{2:2}$-ML297(G1G4) and GIRK1/4$^{2:2}$-ML297(G1G1). a**, Summary table showing the interactions and permeations percentages and their total contribution over the full simulation trajectory. **b-e**, Representative snapshots of dominant interaction pattern with ML297 orientation and key hydrogen-bonding geometry. The percentage indicates the fraction of simulation time this configuration was maintained and the fraction of permeation events that occurred while the system occupied that pattern.

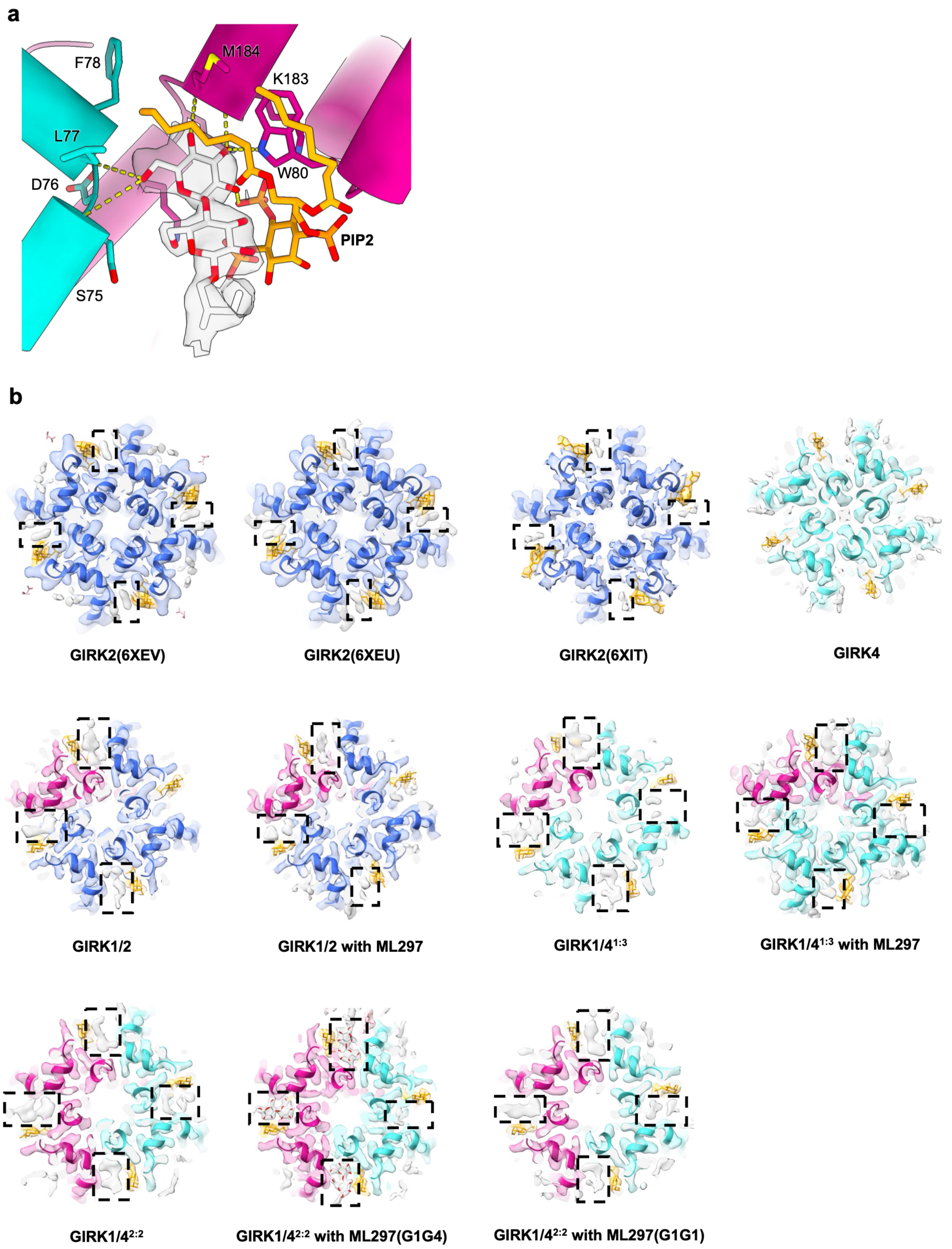


**Extended Data Fig. 16 | Maltoside headgroup of GDN binding near PIP2 binding site.** **a**, A maltoside headgroup from the detergent GDN (white) is positioned adjacent to the pocket, forming hydrogen bonds with $PIP_2$ and nearby residues. **b**, Comparison of previously determined GIRK channel structures (GIRK2: 6XEV, 6XEU, 6XIT; GIRK1/2) and the current GIRK1/4 structures, with and without ML297. In all cases, a similar density adjacent to the $PIP_2$-binding site is observed (boxed).

**Table 1. Cryo-EM data collection, refinement and validation statistics**

| | GIRK1/4$^{1:3}$ | GIRK1/4$^{2:2}$ | GIRK1/4$^{1:3}$–ML297 | GIRK1/4$^{2:2}$–ML297$^{G1G1}$ | GIRK1/4$^{2:2}$–ML297$^{G1G4}$ | GIRK4 |
|---|---|---|---|---|---|---|
| **Data collection and processing** | | | | | | |
| Magnification | 130,000 | 130,000 | 130,000 | 130,000 | 130,000 | 130,000 |
| Voltage (kV) | 300 | 300 | 300 | 300 | 300 | 300 |
| Electron exposure (e–/Å$^2$) | 50 | 50 | 50 | 50 | 50 | 50 |
| Defocus range (μm) | -0.8 to -1.8 | -0.8 to -1.8 | -0.8 to -1.8 | -0.8 to -1.8 | -0.8 to -1.8 | -0.8 to -1.8 |
| Pixel size (Å) | 0.65 | 0.65 | 0.65 | 0.65 | 0.65 | 0.92 |
| Symmetry imposed | C1 | C1 | C1 | C1 | C1 | C4 |
| Initial particle images (no.) | 3,859,854 | 3,859,854 | 4,436,963 | 4,436,963 | 4,436,963 | 327,157 |
| Final particle images (no.) | 91,528 | 44,397 | 25,050 | 22,681 | 92,036 | 7,971 |
| Map resolution (Å) | 2.3 | 2.5 | 2.8 | 2.8 | 2.4 | 2.8 |
| FSC threshold | 0.143 | 0.143 | 0.143 | 0.143 | 0.143 | 0.143 |
| **Refinement** | | | | | | |
| Initial model used (PDB code) | AF-P48549-F1, AF-P48544-F1 | AF-P48549-F1, AF-P48544-F1 | AF-P48549-F1, AF-P48544-F1 | AF-P48549-F1, AF-P48544-F1 | AF-P48549-F1, AF-P48544-F1 | AF-P48544-F1 |
| Map sharpening *B* factor (Å$^2$) | 51.8 | 48.0 | 52.6 | 47.7 | 52.8 | 76.5 |
| Model composition | | | | | | |
| Non-hydrogen atoms | 10851 | 10593 | 10561 | 10602 | 10860 | 10615 |
| Protein residues | 1296 | 1290 | 1296 | 1289 | 1288 | 1300 |
| Ligands | 14 | 13 | 14 | 15 | 33 | 15 |
| Water | 262 | | | | | |
| *B* factors (Å$^2$) | | | | | | |
| Protein | 41.30 | 50.71 | 64.38 | 55.03 | 43.53 | 74.55 |
| Ligand | 52.77 | 69.37 | 88.06 | 74.49 | 56.47 | 95.11 |
| Water | 39.65 | | | | | |
| R.m.s. deviations | | | | | | |
| Bond lengths (Å) | 0.004 | 0.003 | 0.003 | 0.004 | 0.002 | 0.003 |
| Bond angles (°) | 0.929 | 0.574 | 0.656 | 0.919 | 0.564 | 0.556 |
| Validation | | | | | | |
| MolProbity score | 1.36 | 1.64 | 1.61 | 1.88 | 1.37 | 1.95 |
| Clashscore | 3.97 | 4.54 | 3.70 | 5.92 | 4.37 | 8.45 |
| Poor rotamers (%) | 1.7 | 2.68 | 2.62 | 3.05 | 1.61 | 4.46 |
| Ramachandran plot | | | | | | |
| Favored (%) | 98.68 | 97.57 | 97.28 | 96.87 | 98.90 | 99.07 |
| Allowed (%) | 1.32 | 2.43 | 2.72 | 3.13 | 1.10 | 0.93 |
| Disallowed (%) | 0 | 0 | 0 | 0 | 0 | 0 |
